# Molecular simulations reveal the impact of RAMP1 on ligand binding and dynamics of CGRPR heterodimer

**DOI:** 10.1101/2021.05.10.443383

**Authors:** Busecan Aksoydan, Serdar Durdagi

## Abstract

The release of the neuropeptide of calcitonin gene-related peptide (CGRP) plays a key role in the mechanisms of migraine pathology and pain perception as it causes vasodilatation, neurogenic inflammation, mast cell degranulation, sensory signal activation and peripheral sensitivity. Although the findings on the effectiveness of CGRP-targeted therapies in migraine provide new information about the pathophysiology of migraine, questions remain on how the CGRP mechanisms fit into the overall migraine theory. The cryo-EM structure of Gs-protein complexed human CGRP receptor (CGRPR) with bound endogenous CGRP neuropeptide paved the way of understanding the insights into the CGRP receptor function. With several molecular modeling approaches, molecular dynamics (MD) simulations and post-MD analyzes, we aimed to investigate the importance of RAMP1 in the stability of calcitonin receptor-like receptor (CLR). Moreover, we compared the binding modes of the CGRP neuropeptide and CGRPR antagonists (i.e., telcagepant and rimegepant) within the presence or absence of RAMP1. We also investigated the global and local effects of bound molecules on CGRPR as well as their effects on the CLR-RAMP1 interaction interfaces. Results showed that although these molecules stay stable at the ectodomain binding site, they can also bind to the orthosteric ligand binding pocket and form the crucial interactions occurred in the CGRP agonism, which may be interpreted as non-specificity of the ligands, however, most of these interactions at orthosteric site are not sustainable or weak. Particularly, RAMP1 may also be important for the stability of TM domain of CLR hereby stabilizing the orthosteric binding pocket.

## INTRODUCTION

Migraine is a common neurovascular disorder which is characterized by episodes of severe headaches, followed by nausea, vomiting, photophobia, phonophobia and aura (visual impairment) ^1,2^. Although there are broad studies performed to advance our understanding of migraine since the 17^th^ century, the pathophysiology of migraine is still debated, and it is known to be multifactorial ^3,4^. Migraine is generally thought to begin with abnormal activation of the trigeminovascular system, which is responsible for the regulation of blood flow and pain transmission in the head, causing the release of various neuropeptides in the meninges that can cause neurogenic inflammation, such as calcitonin gene-related peptide (CGRP), calcitonin, neurokinin A, and substance P. Inflammation then activates and sensitizes the nerve endings that cause the onset of pain ^5^. CGRP is a calcitonin family member neuropeptide. It is widely expressed and released abundantly from the trigeminal nerves and plays a key role in sensory neurotransmission and pain perception. It causes vasodilatation, sensory signal activation, neurogenic inflammation, mast cell degranulation, and peripheral sensitivity. However, it is protective in hypertensive and inflammatory bowel disease models. Migraines are thought to be the result of a neurogenic inflammation, which then triggers the CGRP release, leading to vasodilation of the central and peripheral nerves and subsequent stimulation of the trigeminal system ^6,7^. The CGRP receptor (CGRPR) is a member of family B GPCRs, is expressed throughout the trigeminal system, including neurons and endothelial cells. Family-B GPCRs contain extracellular, transmembrane (TM), and cytoplasmic domains, as well as accessory proteins such as receptor activity modifying proteins (RAMPs), Na/H exchange regulatory factors (NHERFs), and calmodulin (CaM) ^8^. CGRPR is a heterodimer complex of the calcitonin receptor-like receptor (CLR) and receptor activity-modifying protein 1 (RAMP1) ^9^. RAMPs contain a bulky N-terminal extracellular domain, a single transmembrane (TM) domain, and a short intracellular C terminus. They are essential accessory proteins for the stabilization of the CGRPR complex, particularly for the CLR extracellular domain positioning ^10,11^. Therapeutics for migraine treatment are mostly targeting CLR-RAMP1 protein-protein interaction surfaces, thereby blocking CGRP activity. There are FDA-approved small molecule inhibitors (gepants) and monoclonal antibodies specifically designed to target the CLR-RAMP1 interface rather than CLR-RAMP2 or -RAMP3 ^11,12^. In 2010, the crystal structure of RAMP1-CLR ectodomain complex bound with telcagepant and olcegepant is revealed (PDB: 3N7R ^13^) and it paved the way of understanding how a small molecule inhibitor disrupts the RAMP1-CLR interaction to impair neuropeptide signaling. Moreover, in September 2018, the cryo-EM structure of active, human CGRPR in complexed with Gs protein heterotrimer complex is revealed (PDB: 6E3Y ^14^), and these structures increased the understanding of family-B GPCRs and their functions. Although the findings on the effectiveness of CGRP-targeted therapies in migraine provide new information about the pathophysiology of migraine, questions remain about how CGRP mechanisms fit into the general migraine theory ^6,7^.

Molecular modeling approaches have made significant progress in recent years and in silico studies have become main components of many chemical, physical and biological studies. The protein-protein and protein-ligand interactions are thought to play a role in CGRP biology that are investigated by molecular dynamics (MD) simulations in the current study will contribute to a better understanding of CGRPR biology at atomic level. In this work, we mainly aimed to investigate the following aspects; (i) how the binding of CGRP is affected with and without the presence of RAMP1, (ii) how the CGRPR complex is affected by the binding of a small molecule inhibitor (Rimegepant and Telcagepant), and (iii) what are the differences in dynamical profiles between binding of a gepant molecule to two different binding sites of CGRPR (ectodomain complex interface site and TM domain). The ability to examine these protein-protein and protein-ligand interactions at the atomic level is extremely valuable, both in terms of furthering our understanding of the functions of proteins of clinical and biological importance through directing and guiding experimental studies.

## RESULTS AND DISCUSSION

We divided our systems in two groups for evaluation; (i) CLR-Gs coupled whole structures to investigate CGRP binding and the effects of RAMP1, (ii) CLR complexes without Gs subunits to compare the binding modes of small molecule inhibitors (rimegepant and telcagepant) and the effects of RAMP1 to the binding of these molecules to the receptor. CGRPR has two binding pockets, one in the extracellular CLR-RAMP1 domain (in this work we will refer it to as ***site 1***) and the other one in the CLR TM domain (***site 2***).

### RMSD-time analyzes of CLR-Gs complexes reveal the importance of RAMP1

The initial analysis of 500 ns of MD simulations trajectories indicates that the simulations are converged after around 150 ns of time period (Figure 1). According to these simulations, RAMP1 presence is an important factor for the stabilizing whole structure (CLR - RAMP1 - Gs, Figure 1A, green lines, average root-mean square deviation (RMSD) of 3.9 Å) compared to CLR - Gs complex (pink lines, average RMSD of 7.1 Å. Corresponding average RMSD values of the last 250 ns of simulations for these systems are calculated as 4.6 and 6.5 Å in the given order). However, it can be roughly observed that CGRP binding might not affect the whole structure stability with or without the presence of RAMP1 as we see the timeline (Figure 1A, dark blue, gray and green lines), The average RMSD values are calculated as 4.9 Å and 5.3 Å for CLR-CGRP-Gs and CLR-RAMP1-CGRP-Gs simulations, respectively (the average values of the last 250 ns for the corresponding systems are 5.2 and 5.8 Å, respectively). RMSD-time plots of the replica simulations can be found in Figure S1. According to these basic stability tracks, we can deduce that both gepants may similarly affect CLR-RAMP1 structure when bound to ***site 1***, but rimegepant molecule causes more rigidity to CLR-RAMP1 complex than telcagepant molecule when bound to ***site 2***. Hence, without RAMP1, CLR structure is more flexible with rimegepant compared to telcagepant bound at ***site 2*. RMSD-time analyzes to observe the effects of rimegepant and telcagepant molecules on CLR and CLR-RAMP1 complexes**. Telcagepant is a first generation gepant molecule which has lower bioavailability compared to rimegepant and its development is terminated. Rimegepant, a second generation gepant molecule, is an FDA approved and orally available compound. Since there is no crystal structure of CGRPR complexed with rimegepant, we aimed to compare the co-crystallized structure of telcagepant to our rimegepant-docked complexes. For the gepants bound at ***site 1*** (Figure 1B, dark blue and gray lines), both compounds have similar trends with average RMSD values of 6.0 and 5.7 Å, for rimegepant and telcagepant bound systems, respectively (the average values of last 250 ns RMSD values are 6.6 and 6.3 Å in the given order), which could be interpreted that when bound to the ***site 1***, CLR-RAMP1 heterodimer structure tends to have similar stability trend regardless of bound gepants. Ligand binding ***site 1*** has a common ectodomain binding pocket (ECD) of CLR and RAMP1 interface, which will be further explained in the next sections, it would already be disrupted and there is no necessity to investigate the ligand binding mode at ***site 1*** without RAMP1, therefore we did not perform further simulations for the bound ligands at ***site 1*** without RAMP1. At ***site 2*** (Figure 1B), the case is different than it is at ***site 1***, CLR-RAMP1 complex have different RMSD trends between rimegepant (average RMSD values of 4.7 Å, 4.8 Å in the last 250 ns, pink lines) and telcagepant (average RMSD values of 8.7 Å, 9.4 Å in the last 250 ns, green lines), that may indicate rimegepant is more favorable in ***site 2*** than telcagepant when in complexed with CLR-RAMP1. However, telcagepant binding does not noticeably affect the stability of CLR, regardless of RAMP1 heterodimerization (green and purple lines at Figure 1B, both around 9 Å). Without the presence of RAMP1, rimegepant causes much more flexibility to CLR (cyan lines at Figure 1B, average RMSD values of 12.5 Å, 13.6 Å in the last 250 ns) than telcagepant (magenta lines, average RMSD values of 8.3 Å, 9.4 Å in the last 250 ns). We can also track the translational and rotational motions of these small molecules with respect to protein or the ligand itself considered as reference structures (Lig-fit-Prot and Ligfit-Lig RMSD, respectively) (Figure S2). Here, both gepants keep their internal motions (i.e., LigFitLig) stable at ***site 1*** throughout the simulations (Figure S2A, gray and dark blue lines, both have around 0.5 Å of RMSD values), whereas telcagepant molecule tends to keep its position (average RMSD value of 1.9 Å) in the binding ***site 1*** of CLR-RAMP1 structure more than rimegepant (average RMSD value of 3.8 Å) (Figure S2B). At ***site 2***, we do not observe large fluctuational differences in the internal motions of both gepant molecules (with respect to themselves as reference) regardless of the presence of RAMP1 structure, especially after around 300 ns of MD simulations (Figure S2A). Furthermore, telcagepant molecule also tends to keep its position with respect to protein both with and without the presence of RAMP1 (Figure S2B, green and purple lines) after 200 ns of the MD simulations, whereas rimegepant molecule has large fluctuations and tends to leave the protein without RAMP1 structure (Figure S2B, up to 19 Å, light blue lines), while it has lower RMSD values (around 10 Å, pink lines) and keeps it position after around 150 ns. Corresponding values for replica simulations are given in Figures S3 and S4.

**Figure 1.**
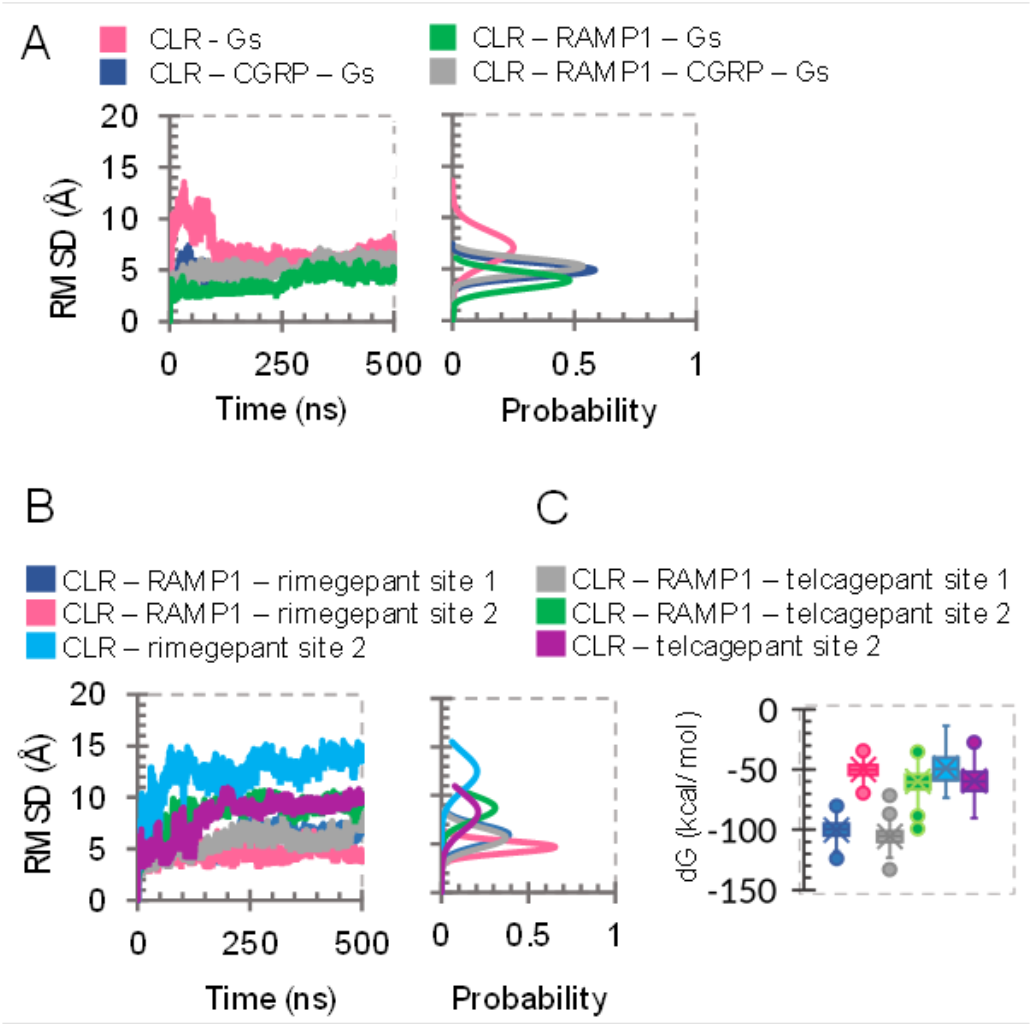
Root-mean-square-deviations vs. simulation time plots of CLR structures in Gs-coupled (A, top) and Gs-uncoupled (B, bottom) complexes. Normal distribution probability plots of RMSD values are plotted next to de corresponding RMSD-time plots, both share the same y axes. Whisker plot of the MM/GBSA dG – NS (non-strain) energy values of the extracted frames of MD simulation trajectories including all replica simulations (C). MMGBSA dG Bind (NS) = Complex – Receptor (from optimized complex) − Ligand (from optimized complex) = MMGBSA dG Bind − Rec Strain − Lig Strain.

### Gepant molecules bind to *site 1* more tightly

We calculated the average MM/GBSA binding free energy values of gepant molecules on both binding sites (Figure 1C). Supporting our Lig-fit-Lig and Lig-fit-Prot data, we observed the most negative dG – NS binding free energies of both gepants (−100.1 and −105.7 kcal/mol for rimegepant and telcagepant structures, respectively) at ***site 1***. However, at ***site 2***, we noted that gepant molecules bind slightly tighter to CLR in the presence of RAMP1 (−50.0 and −60.0 kcal/mol for rimegepant and telcagepant structures, respectively) than to mere CLR structure (−48.9 and −59.6 kcal/mol for the molecules of corresponding order). These calculations are performed for all replicas, and the average of the average dG – NS values are given here. Corresponding values for each simulation replicas and time-based plots are given in Table S1 and Figure S5.

### Global stability of CLR-Gs is severely affected by the absence of RAMP1

We performed principal component analysis (PCA) and plotted the eigenvalue magnitudes for whole protein structures (Figure S6) as well as CLR-only eigenvalues (Figure 2A). In this context, higher eigenvalue magnitude values indicate higher mobility of the structure ^15^. As we compare the CLR-Gs complexes with (gray and green lines at Figure 2A) or without RAMP1 (dark blue and pink lines, Figure 2A), RAMP1 heterodimerization with CLR is observed to be dramatically important in terms of global motion stability. We can see that CGRP binding also lowers the magnitude of CLR motions without RAMP1 structure (dark blue line), yet it is still less stable than bound to CLR in CLR-RAMP1-Gs complex (Figure 2A). Moreover, we also examined the eigenvalue magnitudes of each domains separated in each system (Figure S7) and observed that CLR always has the highest motions compared to the other components (i.e. RAMP1, Gα, Gβ, Gγ and CGRP) followed by Gα subunit. Gβ and Gγ are the most stable ones among the others. Interestingly, CGRP has much lower magnitude of motion when bound to CLR-RAMP1-Gs complex rather than CLR-Gs complex, which may indicate that the stability of CGRP neuropeptide may also depend on the presence of RAMP1 (Figure S7). As we check how these motions affect the topological domains of CLR (Figure 3A) by root mean square fluctuation (RMSF) values, we note the crucial role of RAMP1 structure in the stability of CLR, especially on the extracellular domain. The fluctuations that occurred in CLR-Gs complex (pink lines) are mostly high at extracellular domain, ECL1, ICL2, TM4 and ECL2, particularly. In the CLR-CGRP-Gs system (dark blue lines, Figure 3A), we monitored higher fluctuations in the extracellular domain part, however, as supporting the PCA data given above, the other regions are more stable in the CLR-CGRP-Gs system than in CLR-Gs complex. CLR-RAMP1-CGRP-Gs complex (gray lines, Figure 3A) has the lowest fluctuation trend among all systems, where especially the extracellular, TM4, ECL2 and TM5 domains have the low fluctuations (alpha carbons of the residues are more rigid). Furthermore, we also observed that CGRP neuropeptide bound systems (dark blue and gray lines) have slightly higher fluctuations on helix-8 than CLR-RAMP1-Gs system (green lines, Figure 3A).

**Figure 2.**
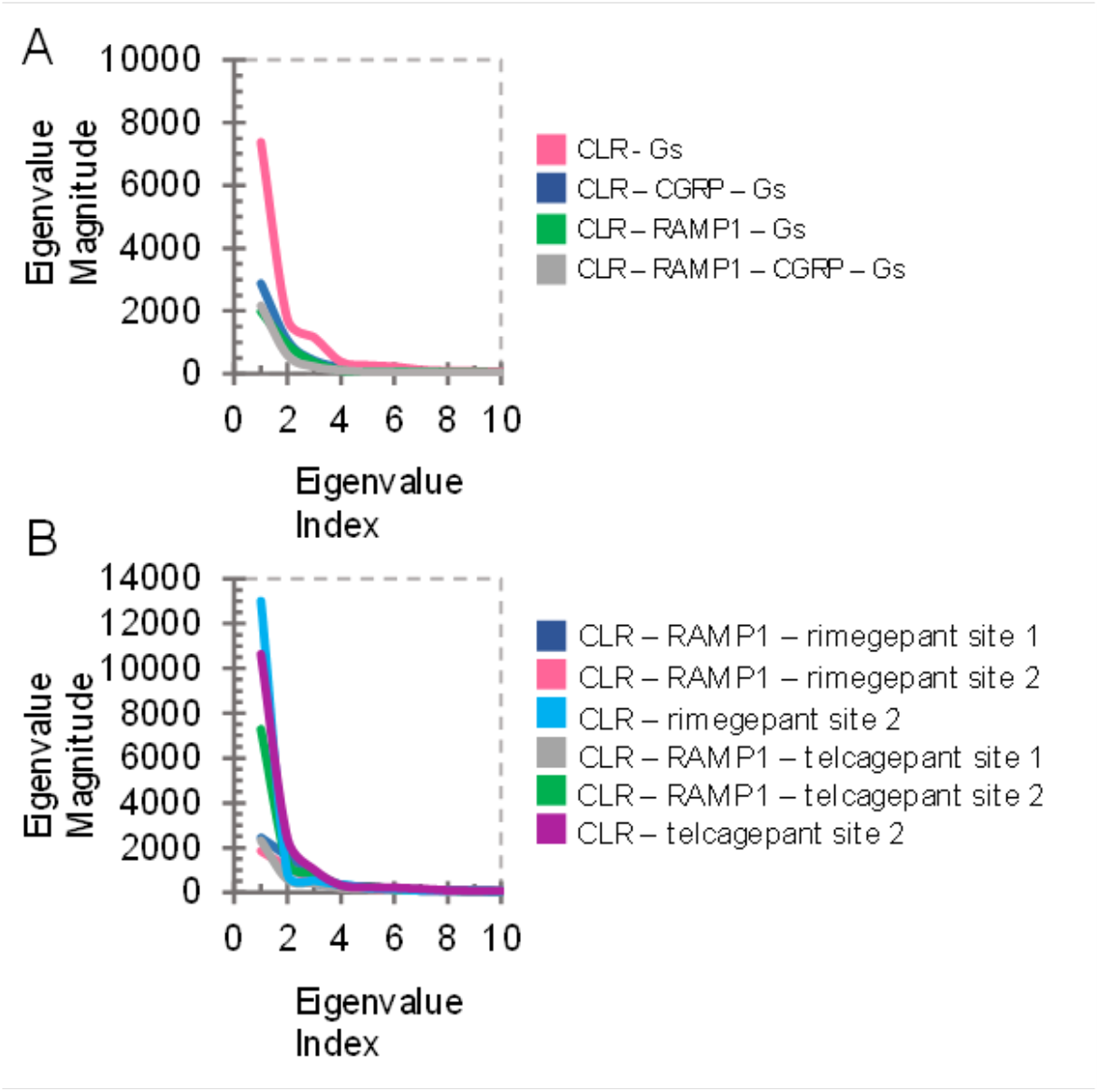
Principal component analysis (PCA) for Gs-c coupled (A, top) and Gs-uncoupled (B, bottom) complexes. Magnitudes of the first ten eigenvalues are plotted. Minor tick marks have a scale of 500 units.

**Figure 3.**
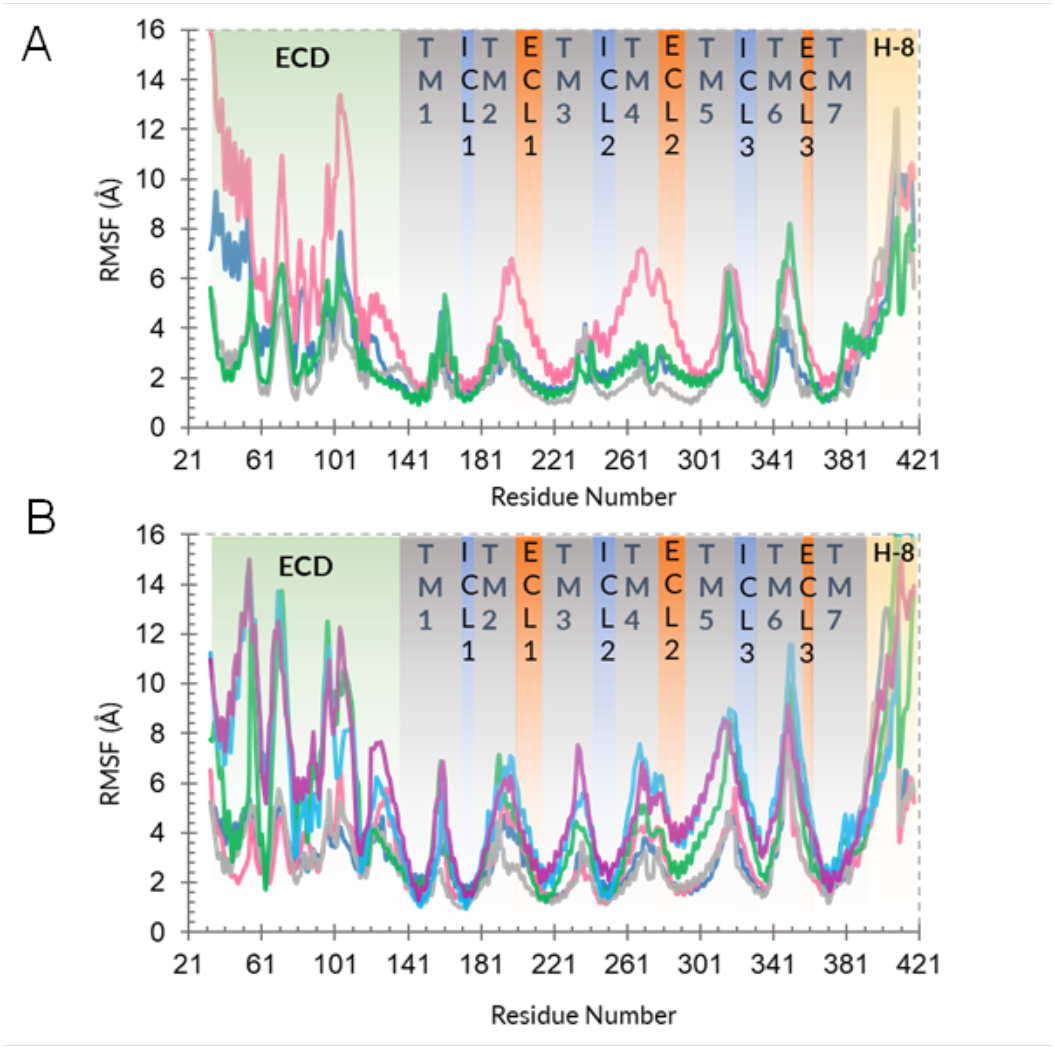
Root-mean-square-fluctuations (RMSF) plots of CLR structures in Gs-coupled (A, top) and Gs-uncoupled (B, bottom) complexes. Topological domains of CLR are divided with colored and labeled columns (ECD: ectodomain, TM1-7: transmembrane domains 1-7, ICL1-3: intracellular loops 1-3, ECL1-3: extracellular loops 1-3, H-8: intracellular helix – 8). A. CLR – Gs: pink lines, CLR – CGRP – Gs: dark blue lines, CLR – RAMP1 – Gs: green lines, CLR – RAMP1 – CGRP – Gs: gray lines. B. CLR – RAMP1 – rimegepant site 1: dark blue lines, CLR – RAMP1 – rimegepant site 2: pink lines, CLR – rimegepant site 2: cyan lines, CLR – RAMP1 – telcagepant site 1: gray lines, CLR – RAMP1 – telcagepant site 2: green lines, CLR – telcagepant site 2: magenta lines

### Telcagepant molecule increases the flexibility in the global structure of CLR-RAMP1 complex compared to rimegepant

The motion magnitudes of CLR in the absence of RAMP1 protein is dramatically larger in all gepant-bound systems, than those that have the RAMP1 heterodimer which indicates the structural instability (Figure 2B). At ***site 1***, rimegepant (dark blue line, Figure 2B) and telcagepant (gray line, Figure 2B), we observed similar effects to the motion magnitudes of CLR-RAMP1 complex. However, at ***site 2***, when bound to rimegepant (pink line, Figure 2B), CLR-RAMP1 complex is much more stable compared to telcagepant (green line, Figure 2B). As supporting our data, we presented in the previous sections, we again observed that without RAMP1, CLR has much more magnitude of motion in rimegepant-bound (cyan) than telcagepant-bound system (magenta). Furthermore, as we examined the component-based magnitude of eigenvalues in each system (Figure S8), the stability of RAMP1 is less affected than CLR upon ligand binding regardless of gepants.

According to our RMSF calculations performed on gepant-bound systems (Figure 3B), CLR-only systems at binding ***site 2*** (CLR-rimegepant and CLR-telcagepant, cyan and magenta lines, respectively) have higher RMSF values at extracellular domain, TMs 3, 4 and 5 among the others. Particularly rimegepant molecule causes the highest fluctuation on TM6 domain residues 346-350, near the ECL3. As we note the rimegepant and telcagepant molecules at ***site 2*** of CLR with RAMP1 (Figure 3B, pink and green lines, respectively), we can interpret that when bound to ***site 2***, CLR is more stable with rimegepant than telcagepant. Telcagepant binding at ***site 2*** seems to affect particularly the residues on extracellular domain and TM5 with higher fluctuations. Supporting our previous PCA and RMSD data, here we also observe that at binding ***site 1*** of CLR-RAMP1 complex, both gepants have lower RMSF values compared to ***site 2*** (rimegepant – dark blue lines, telcagepant – gray lines). Furthermore, as we compared the ***site 1*** versus ***site 2*** RMSF values of rimegepant and telcagepant separately (Figure S9), we observed that both CLR and RAMP1 structures are more stable (having less fluctuations) when bound to rimegepant molecule regardless of binding site, however, telcagepant molecule cause more fluctuations on CLR-RAMP1 complex when bound to binding ***site 2***.

### The changes in cross-correlated motions of CLR by RAMP1 heterodimerization and CGRP binding

We performed residue cross correlation analyses for each simulated system to examine the movement correlation between chosen residue pairs, here we plotted the heatmap for CLR structure (Figure S10) and highlighted regions for CGRP binding and RAMP1 interactions (TM3, TM4, TM5, ECL2 and extracellular domain). Firstly, within the whole picture, we can observe noticeable pattern differences between the systems with and without RAMP1. As we compare CLR-Gs and CLR-RAMP1-Gs systems, the motion correlations are mostly changed in the extracellular domain, followed by TM3 and TM5. For example, the last few residues of TM5 move less correlated and anticorrelated in CLR-Gs than it is in CLR-RAMP1-Gs (Figure S10). We also compared pattern changes when we compared CLR-RAMP1-Gs and CLR-RAMP1-CGRP-Gs systems and observed that the most motion changes occurred in TM5 and TM3, followed by ECL2. Particularly in TM3 (res. 213-240), here we can observe that the motions of the last few residues of TM3 are more correlated in CLR-RAMP1-CGRP-Gs than CLR-RAMP1-Gs. Finally, without RAMP1, we observed much more correlated motions (dense red color) in TM5, where mainly CGRP neuropeptide has interactions; and ECL2, which is also important in CGRP binding (Figure S10). Small molecule inhibitors affect the cross-correlated motion pattern of the CLR extracellular domain more when bound to *site 2*. Although we observed mostly similar effects of rimegepant and telcagepant molecules at binding *site 1* so far, we observed slight changes in the motion correlations of residue pairs of extracellular domains and TM3 of CLR-RAMP1 complex (Figure S11). At *site 1*, most of the neutral motions between residue pairs (white regions) of extracellular domain of CLR for rimegepant-bound complex are observed to move more anticorrelated (blue regions) in telcagepant-bound complex. The same behavior of correlations from correlated to anticorrelated motions are also observed in TM3, the residues in the first half of TM3 move more correlated (dense red color) in complex with rimegepant than with telcagepant (more white and blue regions, Figure S11). At CLR binding ***site 2***, there is a noticeable difference in the patterns between CLR structure with and without RAMP1 (Figure S11). Again, we observed that the most affected region and residue pairs in terms of motion correlation is the extracellular domain of CLR in CLR-RAMP1 complexes with bound gepant molecules. Like at binding ***site 1***, binding of telcagepant molecule to ***site 2*** is observed to cause more anticorrelated motions in the extracellular domain compared to rimegepant molecule which has more correlated motions in the corresponding region. The remaining regions in CLR we outlined for this analysis (TM3, TM4, ECL2 and TM5) do not have many differences regardless of complexed gepants. However, the situation of extracellular domain with the presence of RAMP1 is the opposite in CLR-only systems (Figure S11), where we observed more anticorrelated motions in CLR-rimegepant complex than CLR-telcagepant complex.

Until here, we analyzed the global motion and stability changes on CLR structure with or without RAMP1 accessory protein, as well as the effects of gepant binding on these systems. Overall, according to our MD simulations, it could be deduced that CLR protein stability is severely disrupted without RAMP1 dimerization, regardless of CGRP nor gepant binding. At CLR-RAMP1 ectodomain binding ***site 1***, rimegepant and telcagepant binding cause similar effects on CLR stability, whereas at binding ***site 2***, rimegepant molecule is more stable than telcagepant in terms of translational and rotational movements and particularly in the presence of RAMP1, CLR structure is more stable when in complexed with rimegepant than with telcagepant. From here, we will examine the atomistic interaction details, binding modes of ligand molecules and the effects of RAMP1 protein on these conditions.

### Binding modes of rimegepant and telcagepant at CLR-RAMP1 ectodomain binding *site 1*

As explained in previous sections, we performed our MD simulations for telcagepant-bound CLR-RAMP1 complex at binding ***site 1*** initially based on its co-crystallized conformation in 3N7R coded PDB structure, and we performed molecular docking for rimegepant to obtain the docked complex for MD simulations. We monitored protein interactions with the gepants at the ectodomain complex of CLR-RAMP1 heterodimer.

Telcagepant molecule is observed to keep its crucial interactions ^13^ for antagonism (Figure 4, Figure S12), which are W74 and W84 in RAMP1 ectodomain as well as W72 and T122 in CLR, particularly W74 and W84 residues frame the hydrophobic pocket, thus telcagepant forms and keeps the hydrophobic interactions throughout the simulations. W84 also forms a hydrogen bond from its sidechain to the ligand through a water bridge. At CLR extracellular domain, telcagepant maintains its various interactions (hydrogen bonds, hydrophobic and water bridges) with W72, T120, W121, T122, Y124 and T125 residues. Rimegepant molecule share the same interaction profile with telcagepant but with some extra interactions (Figure 4, Figure S13), with residues such as D71 at RAMP1 and R38, I41 at CLR structure through hydrogen bond, hydrophobic, water bridge and ionic interactions. We also observed that all the crucial interactions mentioned above are maintained throughout the simulations, as well as in the replica simulations.

**Figure 4.**
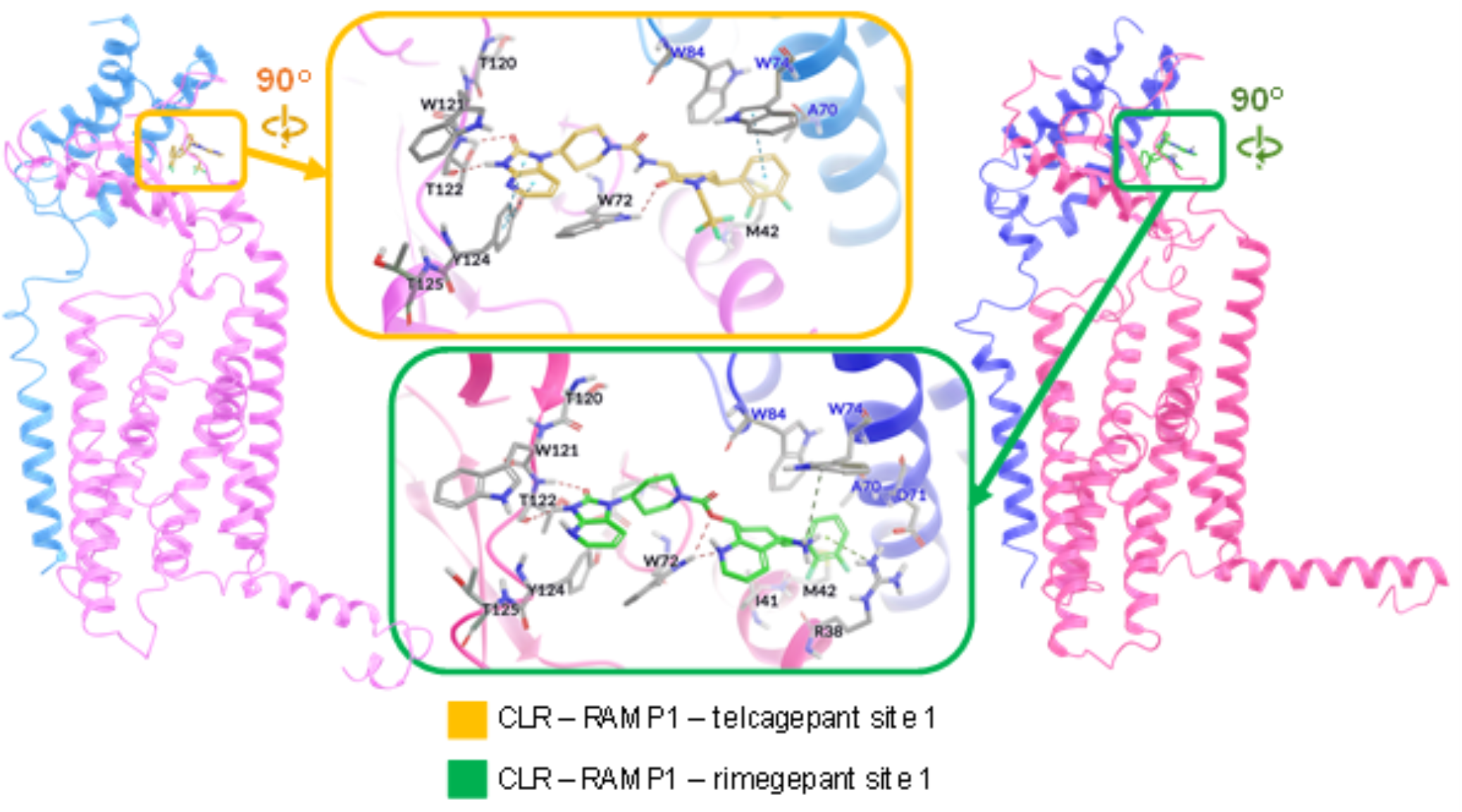
Representative structures of CLR-RAMP1 complexes with bound-gepant molecules at ectodomain binding site (***site 1***). Protein structures are shown in ribbon representation. RAMP1 structures are shown in blue ribbons, CLR structures are shown in pink colors. Zoomed windows display the 90 degrees of rotation in −y axis. In zoomed views, ligand molecules and interacting residues are shown in stick representation. Dashed lines show the H-bonds, salt bridges and pi-pi stacking interactions. The initial position of telcagepant molecule (upper frame, yellow carbons) in CLR binding site is obtained from the co-crystallized structure with PDB code 3N7R, while the initial position of rimegepant molecule (lower frame, green carbons) is obtained from the molecular docking pose with the best docking score & interaction profile. We extracted the representative structures from the concatenated MD simulation trajectories by choosing the one with the closest RMSD to the average structure.

### Changes in the binding modes of CGRP, rimegepant and telcagepant at *site 2* with or without the presence of RAMP1

As it is described in the study of Liang et al. ^14^, there are several interaction pairs of CLR-CGRP essential for the function and receptor signaling: for example, CLR H295 - CGRP T6, CLR H219 - CGRP T9, CLR Y292 - CGRP D3. Furthermore, CLR R274 and W283 are also stated to be important for CGRP signaling. We superimposed the representative structures extracted from our MD simulations of CLR-RAMP1-CGRP and CLR-CGRP systems (Figure 5), it could be roughly observed that there are slight changes in the conformation of CGRP neuropeptide, therefore we plotted the frequency percentages of polar (Figure 5) and non-polar (Table S2) interactions occurred between CLR and CGRP structures in the presence or absence of RAMP1 protein. Firstly, it is seen that most of the interactions occurred from N-and C-termini loops of CGRP are nearly lost (or they are not maintained more than 30% of MD trajectories), such as the interactions between CGRP C2, D3, A5 (N-terminus) – CLR Y292, R355, F349 (TMs 5 and 6) and CGRP T30, G33, S34, F37 (C-terminus) – CLR D94, N128, W121, T122 (extracellular region). As we examined the non-polar interactions, we also observed that most of the interactions occurred in ectodomain region of CLR with CGRP peptide are affected by the absence of RAMP1, where they are not sustained more than 30% of simulation trajectories (Table S2). The main crucial interactions between CGRP T30 – CLR D94 and CGRP F37 – CLR T122 ^13,14^ are observed to be highly persistent with the presence of RAMP1. We also tracked the highly strong CGRP R11 – CLR D366 interactions throughout our MD simulations (Figure 5) as Liang et al. also observed, although they stated that these interactions did not occur in the cryo-EM structure. The CLR H295 - CGRP T6 interactions are stated to be functionally important, which are also highly persistent in our simulations, interestingly, their occurrence is lower in the presence of RAMP1 (70.8%) than the absence of it (79.1%). CGRP T9 and H10 interact with a cluster of residues including T191, L195, H219, I284 and S286 ^14^. Among these, we observed mostly CGRP H10 contacts with CLR H219, I284 and S286, where the polar interactions are persistent regardless of RAMP1 presence (Figure 5), however the nonpolar contacts are disrupted within the presence of RAMP1 (Table S2).

**Figure 5.**
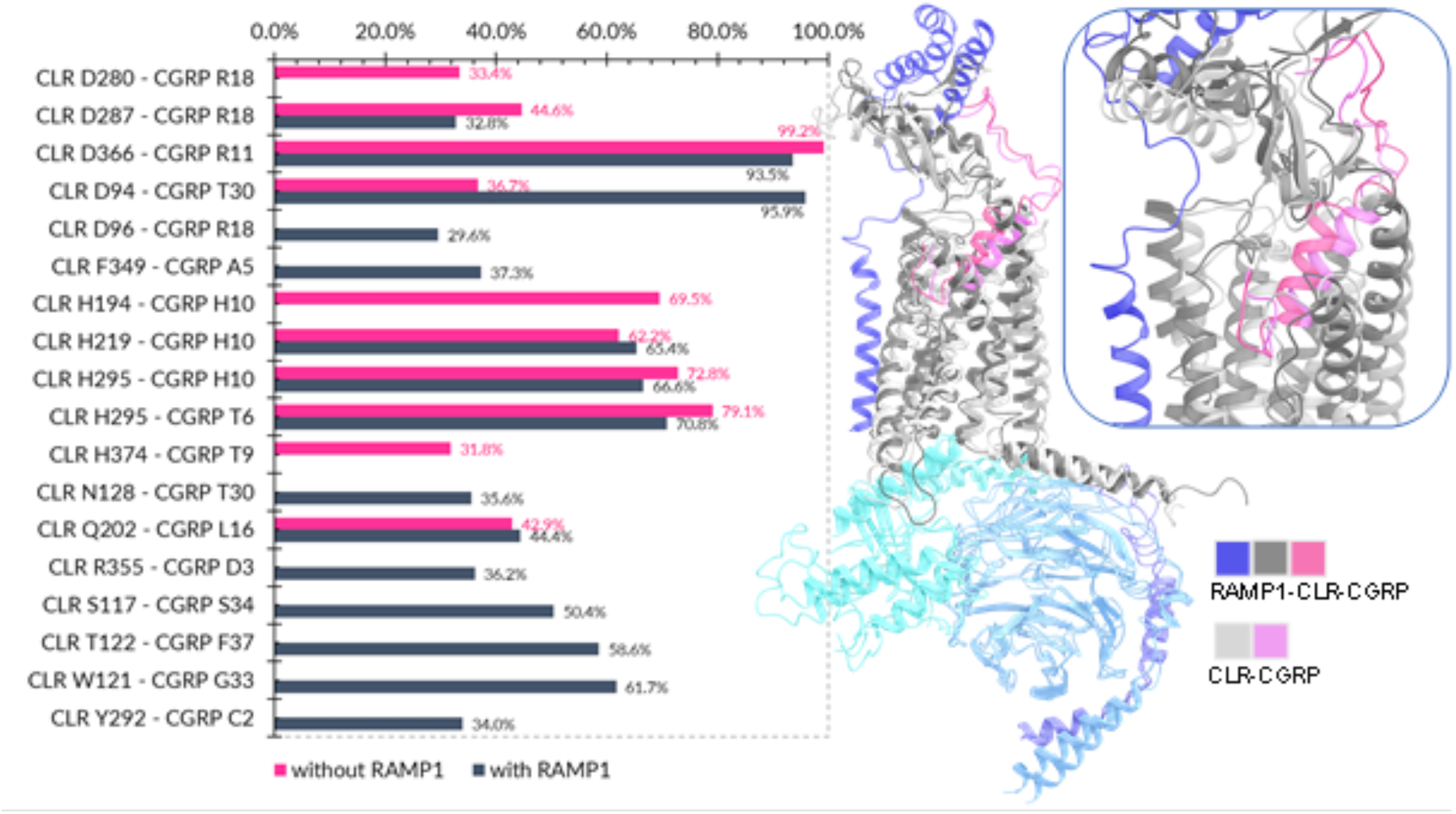
Superimposition of the representative structures of CLR-Gs-CGRP complexes with and without RAMP1 structure (top right corner). Protein structures shown in ribbon representation, G protein subunits of both structures are shown in cyan (Ga), blue (Gb) and purple (Gg) color. RAMP1 structure is shown in dark blue color. CLR structures are represented in dark gray (complexed with RAMP1) and light gray (without RAMP1) color, CGRP neuropeptide is shown as dark and light pink ribbons in the corresponding complexes. Frequency percentages of the polar interactions (H-bonds and salt bridges) occurred between CLR and CGRP peptide with (dark blue bars) and without (pink bars) RAMP1 structure (left). The interactions occurred more than 30% of concatenated MD trajectories are presented.

We also examined the binding modes of telcagepant and rimegepant at ***site 2*** with the presence of RAMP1 (Figures S14-15). At a first glance, we observed that both ligands do not have persistent interactions that occurred less than 30% of simulation time. Both gepants are observed to form hydrogen bonds with H295 through its sidechain, however, these interactions are also weak and could not be maintained throughout the simulations. Here, we will highlight the residue interactions mostly observed to occur in all replicated simulations for each gepant-bound system. For telcagepant molecule (Figure S14), we observed interactions with Y227, H295 and I298 residues that belong to the TM3 and TM5 of CLR and form the base of CGRP neuropeptide binding pocket, as well as F349 among all replicas.

Interactions occurred between telcagepant molecule and Y227 are mostly hydrophobic or water bridges, we also observed hydrogen bond formation through the Y227 sidechain by the O=C–NH linker group of the ligand. H295 sidechain has π-π stacking as well as hydrogen bonding interactions with the oxo-imidazopyridine part of telcagepant. F349 ring also constructs π-π stacking interactions with both rings of oxo-imidazopyridine part, and I298 weakly contributes to the hydrophobic interactions. In the representative structure, we observed that telcagepant has π-π stacking interactions with H295 and F349, and a hydrogen bond with Q376 through its sidechain (Figure S14). Rimegepant shows a different interaction profile compared to telcagepant (Figure S15). Within all replicas, it forms weak water bridges with Y227 similar to telcagepant, however it mostly has contacts with D366, M369, H370, and M373. D366 sidechain forms strong water bridges to the oxygen of the carboxyl linker part of rimegepant molecule. H370 sidechain ring forms π-π stacking interactions with the 6,7,8,9-tetrahydro-5H-cyclohepta[b]pyridine ring and this interaction is maintained throughout the simulations. M369 and M373 residues are also observed to contribute to the common hydrophobic contacts. In the representative structure, interestingly, we captured mostly hydrophobic interactions of rimegepant with M369 and M373 and a hydrogen bond with the backbone carbonyl oxygen of D366 (Figure S15). Without RAMP1, we observed that telcagepant molecule at ***site 2*** has more interactions than rimegepant does, but they are relatively weak and not sustained. Telcagepant molecule has contacts with F184, N187, H219, Y227, H295, M373 and H374, of which the hydrophobic contacts and water bridges through H295 are lost after around 100 ns of simulation time (Figure S16), whereas hydrogen bonds and water bridges with F184 and N187 are formed in the second half of MD simulations. In the representative structure, telcagepant molecule has a conformation that is far from TM3 and TM5, has hydrogen bonds formed with H219, N187, F184 and N226, and it forms π-π stacking interactions with F184 sidechain (Figure S16). Rimegepant molecule has less but stronger interactions throughout MD simulations formed with N187, T191, Y227, Y292, H295, I298, L302 and F349. Among these residues, the hydrophobic and water bridge interactions formed with Y292 and H295 are relatively persistent, however, hydrogen bonds with N187 and T191 could only be maintained around 15% of simulations (Figure S17). Interestingly, in the representative structure of CLR-rimegepant complex at ***site 2***, rimegepant has similar orientation to telcagepant with hydrogen bonds formed with N187, T191 and H219 (Figure S17) whereas H295 does not form any contacts to the ligand.

### Changes in the interactions of CLR-RAMP1 domain interface upon CGRP, rimegepant and telcagepant binding to sites 1 and 2

We tracked all types of polar and nonpolar interactions that occurred between CLR and RAMP1 domain interfaces including both membrane and ectodomain regions (Tables 1 and S3). CLR-RAMP1 ectodomain interface has lots of contacts, most of them are known to stabilize CLR extracellular region, also RAMP1 membrane-embedded helix interacts mostly with TM3, TM4 and TM5 of CLR ^14^. According to our simulations, we observed that the persistency (frequency) pattern of the residue contacts (both electrostatic and hydrophobic) changes in different conditions, i.e. with bound ligand or the binding sites. In extracellular region, numerous interactions that stabilize the heterodimer predominantly occur between CLR helix residues (res. 34-54) and RAMP1 helices (res. 56-85 and 86-101) with C-terminal loop (res. 102-115) ^13,14^. For example, CLR E47 – RAMP1 R112 and CLR ECL2 T288/H289 – RAMP1 D113 interactions are stated to be important because these contacts help to stabilize ECL2 and to link the extracellular and TM domain of RAMP1. According to our simulations (Table 1), these interactions occurred and maintained throughout the simulations of the CLR-RAMP1-Gs system. However, in the CLR-RAMP1-CGRP-Gs system, we observed that the contacts of CLR ECL2 T288/H289 – RAMP1 D113 are dramatically lost from more than 95% to below 40%, yet CLR E47 – RAMP1 R112 is maintained around 60% of simulations. Interestingly, regardless of binding site, rimegepant binding may cause the frequency of CLR E47 – RAMP1 R112 to be lower than 40%, opposite to telcagepant, which nearly did not affect this interaction (keeping the frequency around 50%). On the other hand, telcagepant binding is observed to affect particularly CLR ECL2 T288– RAMP1 D113 interaction more than rimegepant, lowering the frequency from more than 95% to less than 55%. Another ECD interaction pairs, CLR E47 – RAMP1 S107/G108 are affected particularly in rimegepant-bound system at ***site 1***, where their contact occurrence frequency is again lowered to less than 40%. In telcagepant bound systems (both binding sites), the interactions between E47 – G108 pair is more affected than E47 – S107 interactions. The other contacts occurred in ECD interface, between CLR Q50, R38, Y49 and RAMP1 H97, D71 and D90 are not observed to be severely affected, regardless of bound molecule nor binding site (Table 1). We also observed changes and loss in some of the contacts between RAMP1 membrane helix and CLR TMs 3,4 and 5 (Tables 1 and S3). One of the important interactions occurred between RAMP1 membrane helix and CLR TM4 is the Y255 of CLR – S141 of RAMP1 which maintained 78% of total trajectories. These contacts are disrupted, and their occurrence frequency are lowered down to 40% and below in CGRP-bound, rimegepant-bound (both binding sites) and telcagepant-bound (binding ***site 2***) systems. Similar pattern is also observed between CLR I235 (TM3) – RAMP1 V138 residue pair. Other contacts occurred between CLR F262 (TM4) and RAMP1 V133 is affected by gepant binding, where these contacts are observed to persist more than the half of the total trajectories in rimegepant-bound systems, but they are mostly lost in more than 60% of simulations in telcagepant-bound systems. We did not observe any visible changes in the contacts between CLR TM5 and RAMP1 residues (Table S3). Overall, according to our simulation data, it can be interpreted that most of the interactions occurred in CLR-RAMP1 ectodomain interface (CLR ECD, ECL2 – RAMP1 ECD) are affected by ligand binding more than the membrane interface (CLR TMs 3,4 and 5 – RAMP1 membrane helix).

**Table 1.** Frequency of all polar and non-polar contacts occurred CLR-RAMP1 interface.

| Residue Pairs | Gs | Gs +<br>CGRP | Rime-<br>site1 | Rime-<br>site2 | Telca-<br>site1 | Telca-<br>site2 |
| --- | --- | --- | --- | --- | --- | --- |
| CLR E47 - RAMP1 G108 | 53.2% | n/a | n/a | 57.3% | n/a | n/a |
| CLR E47 - RAMP1 R112 | 49.5% | 56.6% | n/a | n/a | 44.0% | 50.4% |
| CLR E47 - RAMP1 S107 | 47.3% | n/a | n/a | 59.4% | 44.0% | 43.8% |
| CLR M42 - RAMP1 A70 | 63.7% | 63.1% | 99.2% | 73.5% | 99.2% | 63.1% |
| CLR M42 - RAMP1 R67 | 75.6% | 77.7% | 58.7% | 67.5% | 51.5% | 74.9% |
| CLR M42 - RAMP1 W59 | 79.3% | 74.1% | 71.3% | 68.1% | 75.8% | 65.3% |
| CLR M42 - RAMP1 Y66 | 52.8% | 51.0% | 55.3% | 58.0% | 76.2% | 54.9% |
| CLR M53 - RAMP1 G98 | 56.7% | 58.4% | 53.0% | 68.4% | 48.1% | 56.7% |
| CLR M53 - RAMP1 H97 | 64.2% | 66.6% | 66.1% | 72.0% | 62.1% | 69.3% |
| CLR M53 - RAMP1 L94 | 62.5% | 56.2% | 56.3% | 52.8% | 43.2% | 50.7% |
| CLR M53 - RAMP1 R102 | 58.7% | 57.1% | 59.3% | 60.4% | 57.7% | 53.8% |
| CLR Q50 - RAMP1 C104 | 74.7% | 82.9% | 66.7% | 61.5% | 81.4% | 69.0% |
| CLR Q50 - RAMP1 H97 | 99.3% | 97.6% | 98.2% | 97.1% | 97.4% | 90.4% |
| CLR Q50 - RAMP1 I106 | 66.3% | 66.5% | 65.0% | 77.6% | 71.0% | 72.8% |
| CLR R38 - RAMP1 D71 | 93.9% | 96.6% | 96.2% | 82.1% | 94.1% | 98.7% |
| CLR Y46 - RAMP1 W59 | 99.3% | 96.5% | 99.0% | 99.1% | 99.7% | 97.4% |
| CLR Y46 - RAMP1 Y66 | 70.9% | 76.1% | 68.3% | 66.3% | 60.2% | 64.3% |
| CLR Y49 - RAMP1 D90 | 97.1% | 98.0% | 99.8% | 92.1% | 96.2% | 88.0% |
| CLR Y49 - RAMP1 F93 | 92.8% | 96.6% | 93.5% | 88.4% | 89.7% | 90.0% |
| CLR Y49 - RAMP1 H97 | 86.3% | 86.7% | 84.4% | 91.7% | 90.5% | 92.0% |
| CLR Y49 - RAMP1 L94 | 42.7% | 52.2% | 45.8% | n/a | 52.3% | n/a |
| CLR H289 - RAMP1 D113 | 99.6% | n/a | 98.8% | 90.4% | 59.7% | 86.4% |
| CLR H289 - RAMP1 L119 | 72.7% | n/a | 62.2% | 68.3% | 61.8% | 55.0% |
| CLR T288 - RAMP1 D113 | 97.1% | n/a | 91.6% | 86.8% | 52.3% | 40.5% |
| CLR Y278 - RAMP1 R112 | 49.2% | n/a | 94.5% | 58.5% | 51.6% | 72.4% |
| CLR I235 - RAMP1 V138 | 47.7% | n/a | n/a | n/a | 51.4% | n/a |
| CLR L231 - RAMP1 T134 | 61.5% | 71.0% | 63.3% | 69.0% | 71.7% | 67.2% |
| CLR T239 - RAMP1 S141 | 69.2% | 59.5% | 46.4% | n/a | 49.6% | 40.0% |
| CLR F262 - RAMP1 T130 | 85.4% | 72.5% | 75.0% | 80.8% | 83.8% | 82.5% |
| CLR F262 - RAMP1 V133 | 45.0% | 41.5% | 50.6% | 56.7% | n/a | n/a |
| CLR L258 - RAMP1 V133 | 54.8% | 53.6% | 57.8% | 45.4% | 67.0% | 60.7% |
| CLR W254 - RAMP1 Q140 | 84.5% | 59.1% | 87.8% | 86.8% | 87.8% | 76.5% |
| CLR Y255 - RAMP1 S141 | 78.1% | 41.0% | n/a | n/a | 61.2% | n/a |

Interactions occurred more than 40 percent of concatenated MD trajectories are presented. The cells with n/a represent the interactions occurred less than 40 percent of concatenated MD trajectories. Rime-: Rimegepant, Telca-: Telcagepant

Molecular modeling and simulation methods have become a standard tool to visualize and mimic various physiological systems and to study cellular phenomena in multidimensional approaches. Till date, the role of RAMPs in class B GPCRs are broadly investigated ^11,16–19^, and there are various experimental studies performed for the neuropeptide-cell signaling and activation mechanisms ^16,19–22^, as well as the receptor stability and ligand selectivity ^9,23,24^. The release of whole structural domains of the human CGRP receptor (CLR + RAMP1) with bound CGRP neuropeptide and in complex with the Gs-protein heterotrimer by Liang et al. is expected to give great momentum to the in silico studies in this area. RAMP1 is mostly studied for its role in the CGRP signaling, where the activation of CGRP receptor predominantly results in vasodilation which is shown in animal models ^25^ and also known to be clinically linked to the migraine pathophysiology ^26^. The therapeutic targeting of CGRP receptor is mostly carried out at the CLR-RAMP1 ectodomain interface, where the C-terminal loop of CGRP neuropeptide is bound, hereby blocking the CGRP binding and signaling which may lead to the decreased CGRP activity for the treatment of migraine. Currently, CGRP activity blockade is being studied by using either monocolonal antibodies or the small molecule inhibitors (gepants). Erenumab is the first monocolonal antibody which is FDA-approved in 2018 for the treatment of chronic migraine ^27^, and rimegepant is a second generation gepant and is a small molecule CGRP receptor inhibitor (approved by FDA in 2020) for the treatment of an acute migraine in adults ^28^. Various gepant molecules are being studied for years and their effects on patients with migraine are broadly reviewed ^29–31^. Both types of molecules are targeting the CGRP receptor to disrupt CLR-RAMP1 interface interactions upon CGRP neuropeptide binding. Here, in this work we aimed to demonstrate firstly how CLR is globally and locally affected by the presence or absence of RAMP1 in the conditions of Gs-coupled (apo and CGRP-bound) systems. Second, we investigated how ligand binding affects the CLR via gepant-bound systems (telcagepant and rimegepant bound without Gs) in ectodomain site (***site 1***) and orthosteric binding site (***site 2***), as well as how these systems are affected with the presence or absence of RAMP1. Thus, we aimed to investigate the role of RAMP1 on CLR and ligand binding, as well as to examine the binding modes and differences of small molecule inhibitors. Moreover, we tracked all the interactions occurred between the CLR-RAMP1 heterodimerization interface, including the extracellular and transmembrane domains, to investigate the effects of CGRP and gepant binding to these domain interactions.

## METHODS

### Modeling and Structure Preparation

The cryo-EM structure of active CGRPR with Gs heterotrimer subunits complex is used to build our systems (PDB, 6E3Y). There are missing loops and residues ^14^ especially in CGRP neuropeptide and CLR parts. We used “Crosslink Proteins” module of Schrodinger’s Crosslink Proteins module ^32^ to add missing residues in the neuropeptide, with the UNIPROT FASTA sequence as a guide ^33^, and GPCRDB ^34^ to obtain a filled and refined model of CLR. In addition, the nanobody structure (Nb35) is removed before the preparation processes. Protein Preparation Wizard (PrepWiz) module of Maestro ^35^ is used to obtain refined, optimized and minimized structures for further simulations. Hydrogens were added, disulfide bonds were formed, missing sidechains were filled ^36^. PROPKA ^37^ is used for the prediction of protonation states of the residues in physiological pH (pH 7.4). A restrained minimization is applied afterwards until the convergence of heavy atoms to 0.3 Å RMSD in OPLS3 forcefield ^38^. CGRPR has two binding pockets, one in the extracellular CLR-RAMP1 domain (***site 1***) and the other one in the CLR transmembrane domain (***site 2***). For the systems with small molecule inhibitors we used two approaches to build our structures; (i) merge the co-crystallized ligand conformation, and (ii) molecular docking. **Ligand Preparation and Molecular Docking**.Telcagepant and rimegepant molecules are used to build small-molecule complexed systems. We used the crystal structure of telcagepant bound ectodomain complex (PDB: 3N7R) to get the ligand 3D structure as well as its bound conformation in ***site 1***. The remaining complexes of these molecules in sites 1 and 2 are generated via molecular docking. The 3D structure of rimegepant is obtained from the PubChem database. LigPrep module is used to prepare the ligand states in pH 7.4 with Epik ^39,40^, and a minimization step is performed to get the energetically favorable ligand structures for molecular docking. Glide module ^41^ of Schrodinger is used to perform grid generation and molecular docking. Grid center for the ***site 1*** is considered as the co-crystallized telcagepant coordinates, while for the ***site 2***; S286, Y292, I293 and H295 residues, which are known as the crucial interactions formed between CLR and CGRP, are centered. Standard Precision (SP) protocol is used for docking and ligands are treated as flexible. Overall, we prepared several systems, including apo-state, with or without presence of RAMP1, and Gs subunits for MD simulations (Table S4). **Molecular Dynamics (MD) Simulations**. Desmond module ^42^ of Schrodinger is used to generate system boxes and perform MD simulations. All the systems are prepared considering OPLS3 forcefield. Each system box consists of POPC bilayer membrane, the membrane positioning coordinates are obtained from OPM server (Orientations of Proteins in Membranes ^43^). The systems are then solvated with TIP3P water model, 0.15 M concentration of NaCl ions and enough amounts of Cl ions to neutralize total charge of the system. The systems are simulated for 500 ns of production (and 1 replica each) with 10000 recorded frames per simulation. NPgT ensemble is used, with constant 310K temperature (Nose-Hoover thermostat ^44^) and 1.01 bar of pressure (Martyna-Tobias-Klein barostat ^45^), as well as 4000 bar.Å of surface tension applied to the membrane bilayer. An extra membrane relaxation protocol is applied before production simulations. Schrodinger’s Maestro, VMD ^46^, and Bio3D package for R ^15^ programs are used to further analysis and visualization of the MD trajectories. **Post-MD Analyses and MM/GBSA Calculations**. We refined the sequence intervals for the topology of each region (extracellular, TMs and loops) based on the POPC lipid thickness in our systems and membrane orientation coordinates (Table S5). In total, we have performed 11.5 µs of MD simulations for 10 different systems given in (Table S4) considering their replicas, and merged all the trajectory replicas for each system (i.e. 20000 or 30000 frames in total for each system) to perform root-mean-square fluctuations (RMSF), principal component analysis (PCA) and cross-correlation analyses. We also calculated root-mean-square-deviations (RMSD) for protein backbone, and for ligand molecules (Lig-fit-Prot and Lig-fit-Lig). ‘Lig fit Prot’ shows the RMSD of a ligand when the protein-ligand complex is first aligned on the protein backbone of the reference and then the RMSD of the ligand heavy atoms is measured. If the values observed are significantly larger than the RMSD of the protein, then it is likely that the ligand has diffused away from its initial binding site. ‘Lig fit Lig’ shows the RMSD of a ligand that is aligned and measured just on its reference conformation. This RMSD value measures the internal fluctuations of the ligand atoms. Moreover, we also plotted protein-ligand contacts histograms and diagrams to monitor the protein interactions with the ligand throughout the simulations. Binding free energy prediction of gepant-bound systems throughout the simulations were performed via Prime-MM/GBSA (Molecular Mechanics / Generalized-Born Surface Area) module. Protein-ligand complexes were extracted from trajectories and solvated using VSGB 2.0 solvation model ^47^, under OPLS3 forcefield. We extracted representative structures for the static analyses from the concatenated MD simulation trajectories by choosing the one with the closest RMSD to the average structure. In addition, we used getcontacts python script (https://getcontacts.github.io/) to calculate frequencies of the interactions of all types (i.e. hydrogen bond, π-cation, π-π stacking, salt bridge, VdW, etc) occurred in CLR-RAMP1 interface within corresponding simulations, as well as to calculate and track the interactions between CGRP neuropeptide and CLR receptor with and without presence of RAMP1 protein.

## ACKNOWLEDGMENT

The numerical calculations reported in this paper were partially performed at TUBITAK ULAKBIM, High Performance and Grid Computing Center (TRUBA resources).

## Notes

### Competing Interest Statement

The authors have declared no competing interest.

