## Supporting Information for "Molecular simulations reveal the impact of RAMP1 on ligand binding and dynamics of CGRPR heterodimer"

### Supplementary Figures

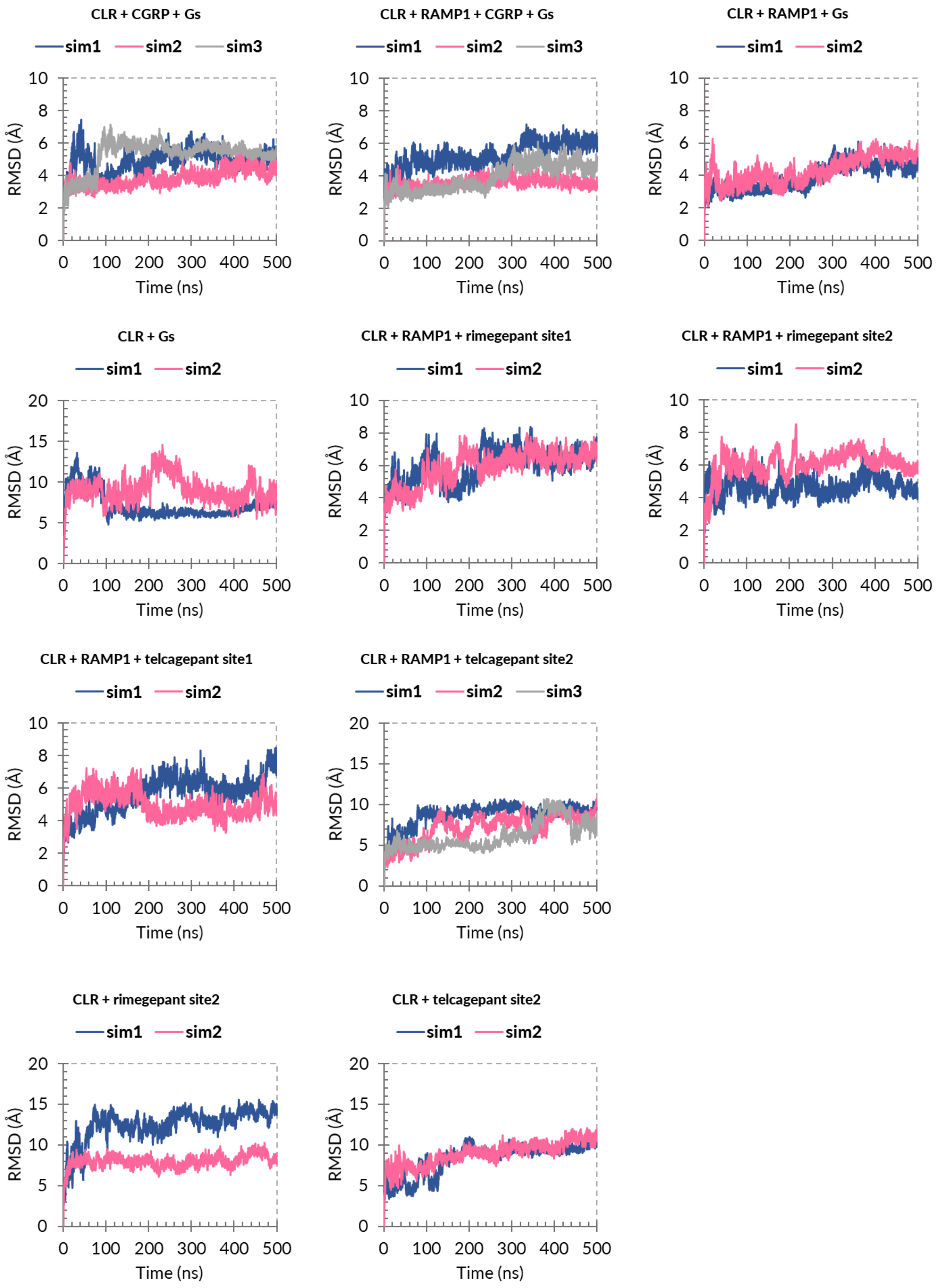


**Figure S1.** Whole protein root-mean-square-deviations (RMSD) vs. simulation time plots of CLR structures in Gs-coupled and Gs-uncoupled systems with corresponding replica simulations.

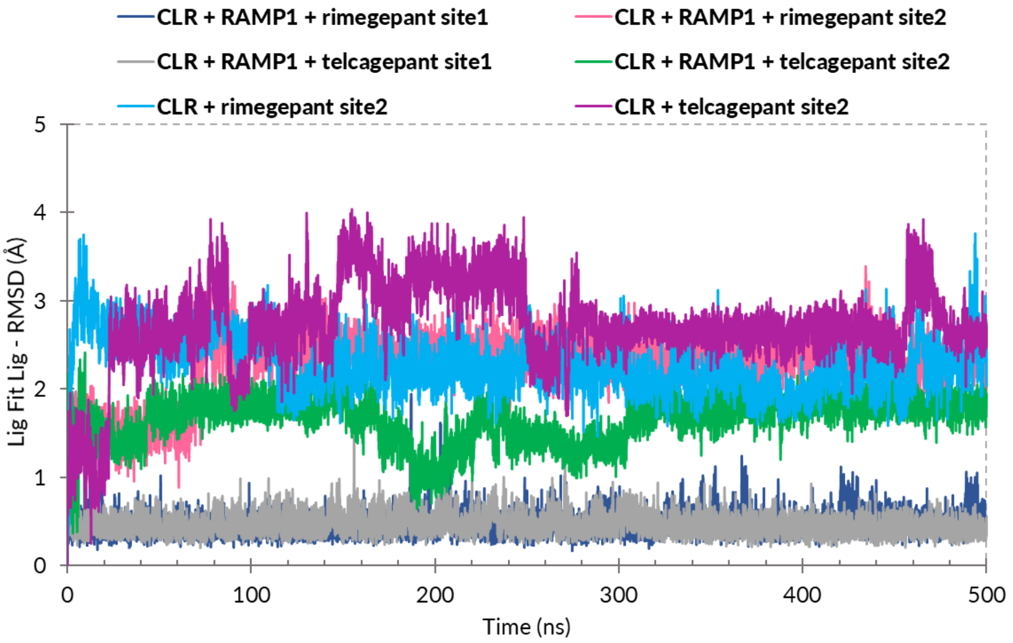


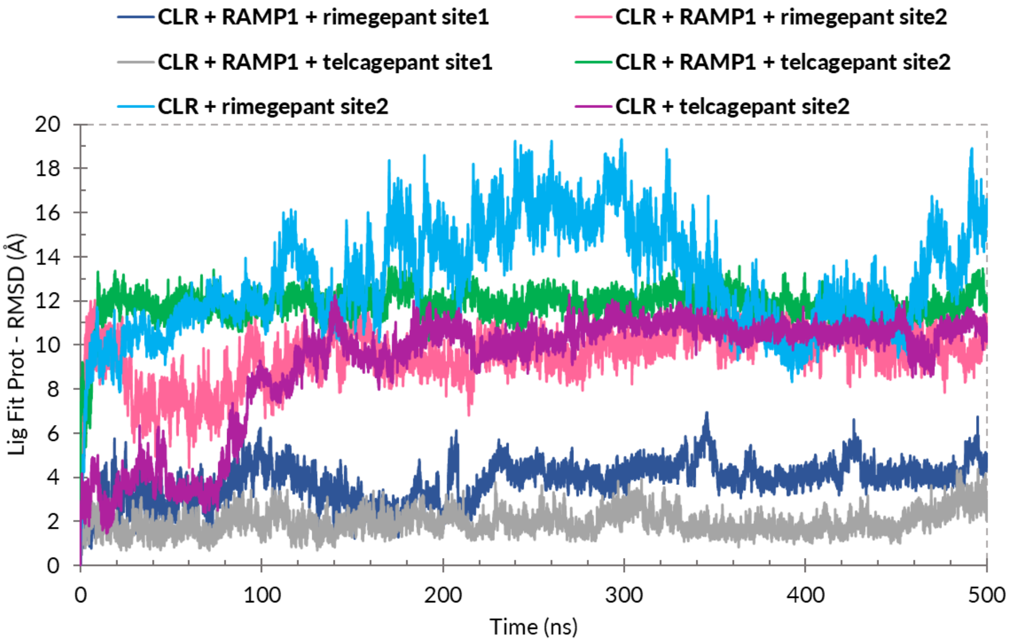


**Figure S2.** Lig-fit-Lig (top; the changes in the atomic positions of the ligand molecule with respect to itself as initial reference position) and Lig-fit-Prot (bottom; the changes in the atomic positions of the ligand molecule with respect to protein as initial reference position) RMSD vs. time plots for Gs-coupled and Gs-uncoupled systems.

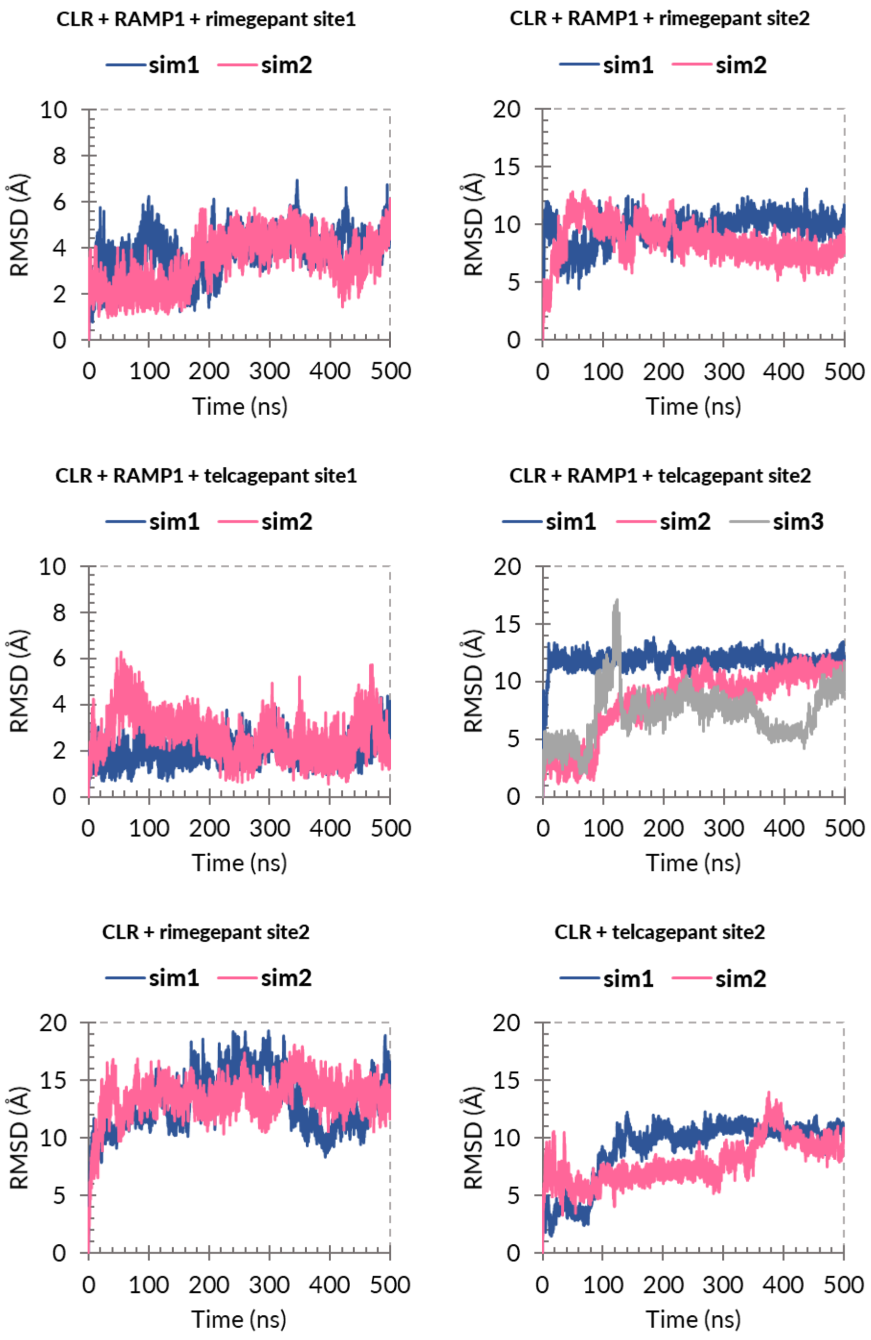


**Figure S3.** Lig-fit-Prot (the changes in the atomic positions of the ligand molecule with respect to protein as initial reference position) RMSD vs. time plots for Gs-coupled and Gs-uncoupled systems with corresponding replica simulations.

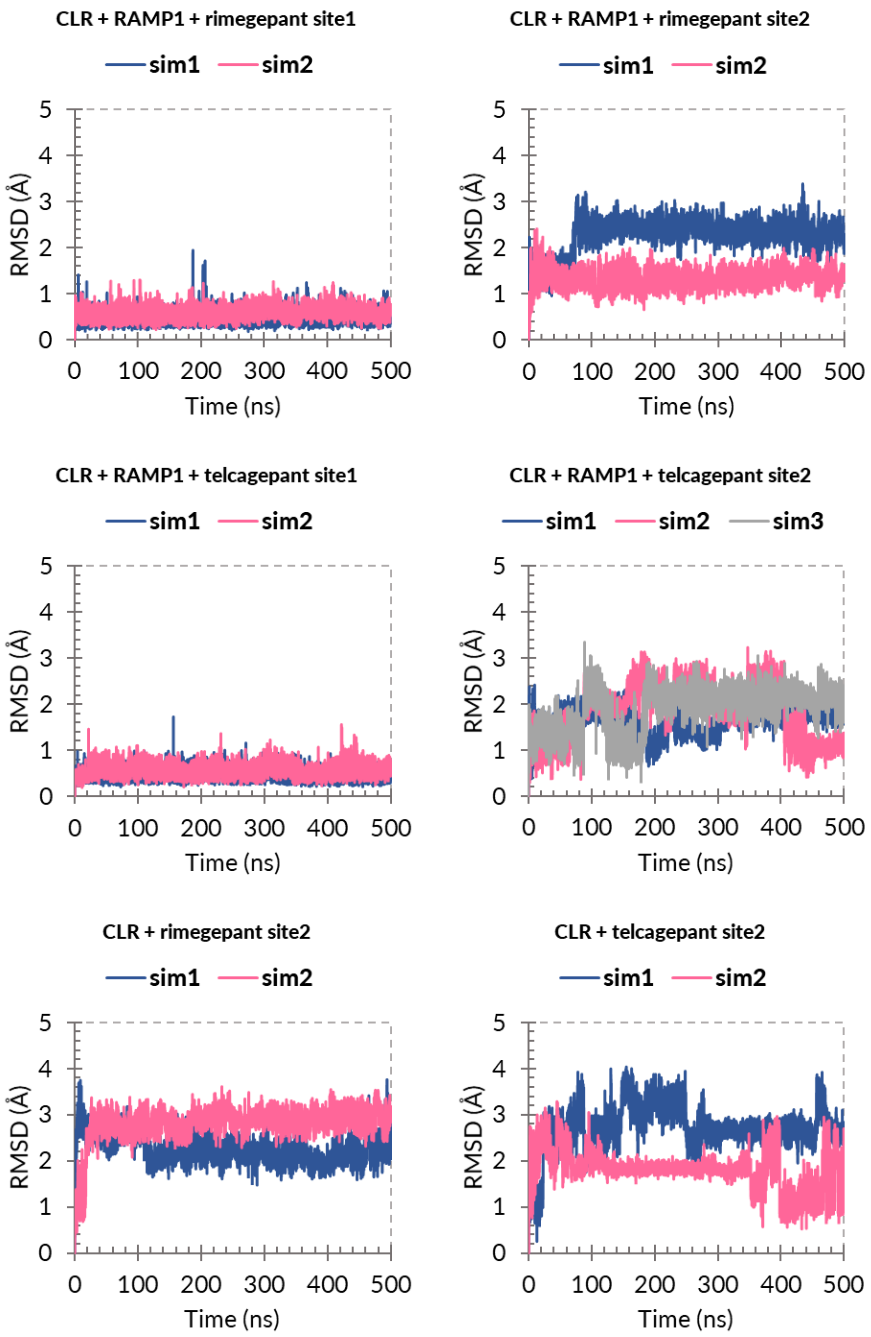


**Figure S4.** Lig-fit-Lig (the changes in the atomic positions of the ligand molecule with respect to itself as initial reference position) RMSD vs. time plots for Gs-coupled and Gs-uncoupled systems with corresponding replica simulations.

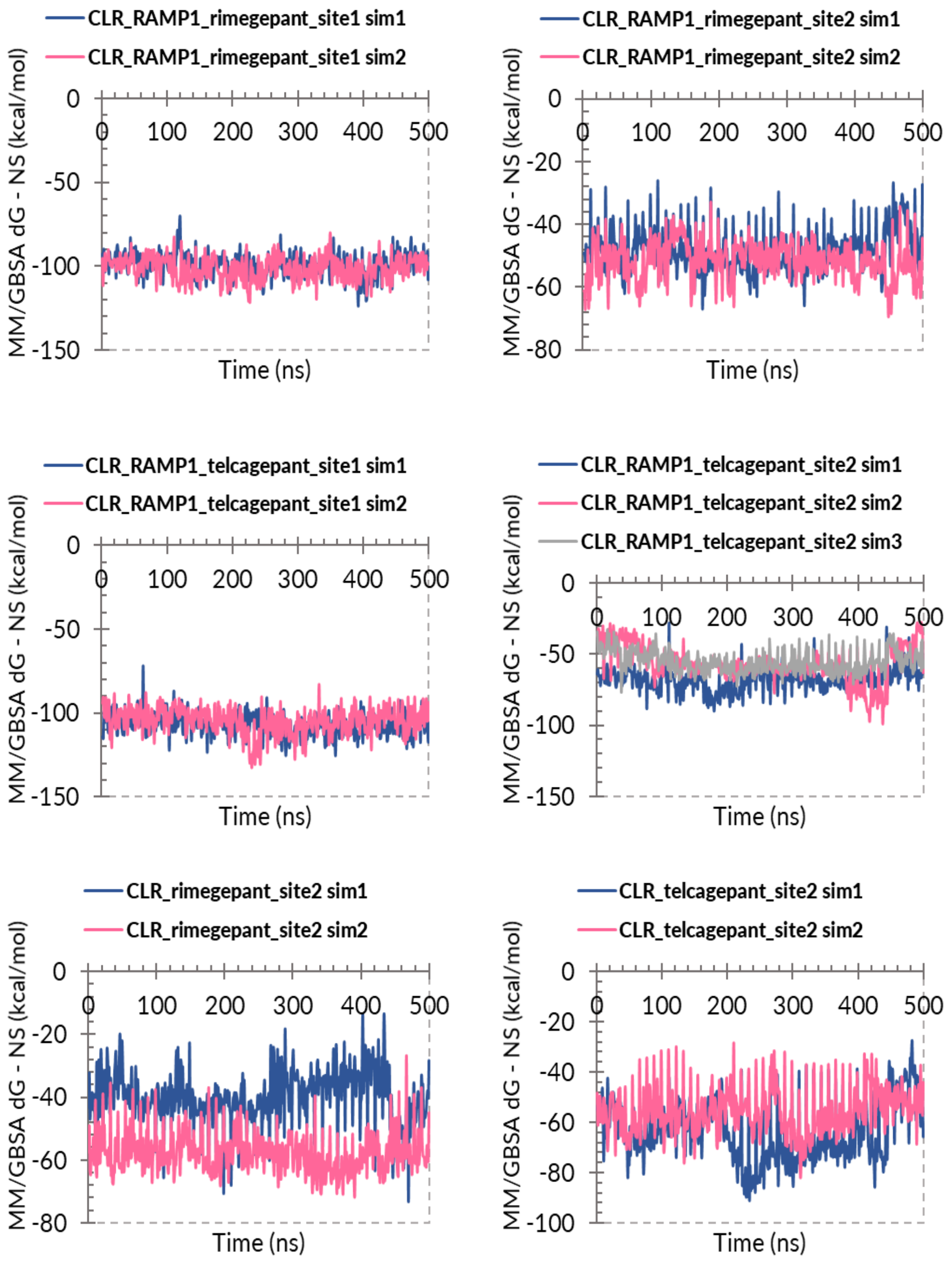


**Figure S5.** MM/GBSA dG-NS (nonstrain) energy vs time plots of the extracted frames of MD simulation trajectories including all replica simulations. MMGBSA dG Bind(NS) = Complex - Receptor(from optimized complex) - Ligand(from optimized complex) = MMGBSA dG Bind - Rec Strain – Lig Strain.

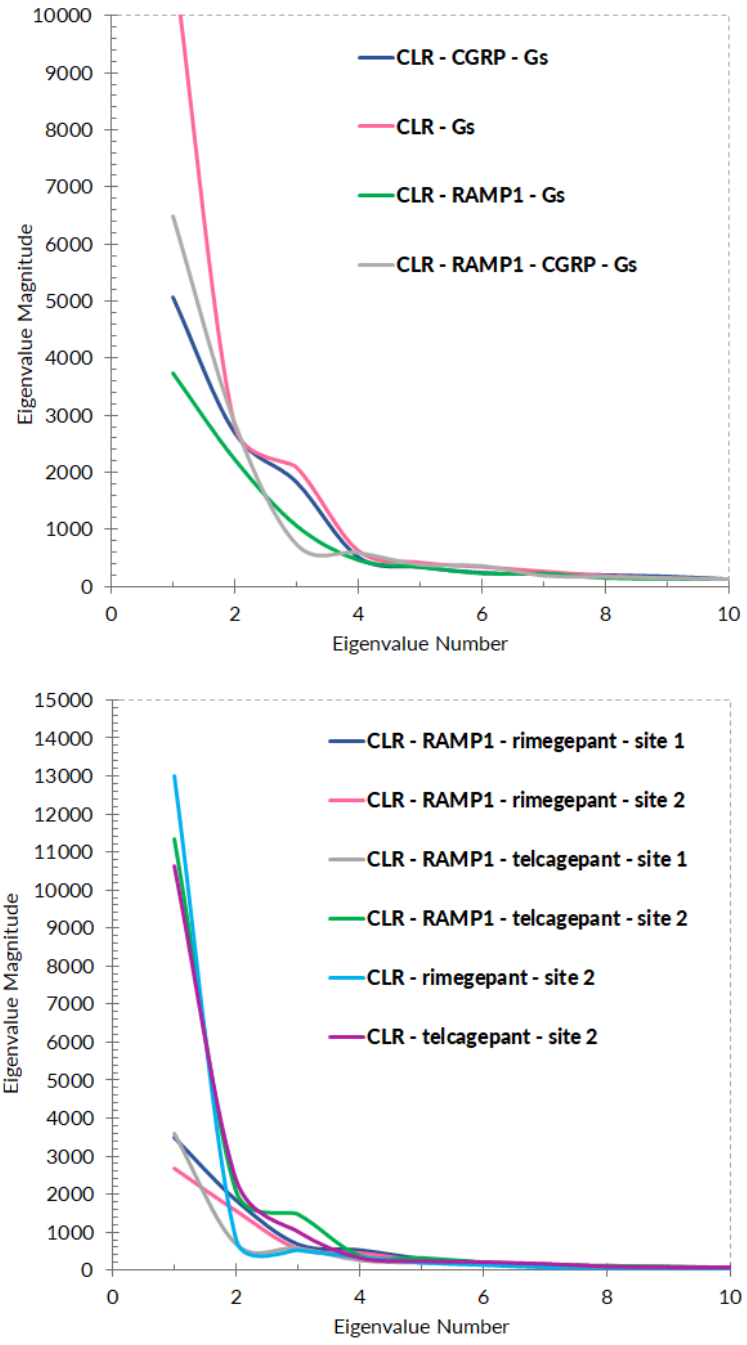


**Figure S6.** Principal component analysis (PCA) for Gs-coupled (A, top) and Gs-uncoupled (B, bottom) whole complexes. Magnitudes of the first ten eigenvalues are plotted.

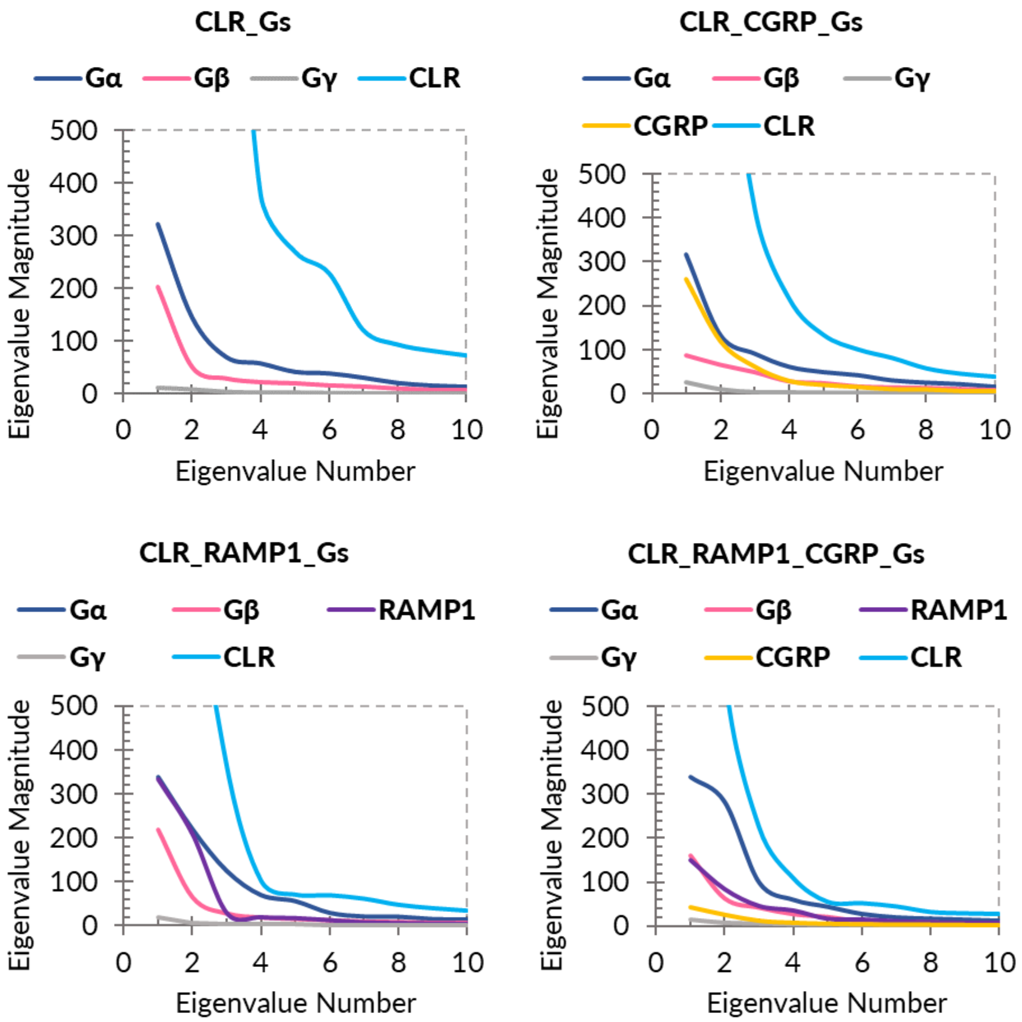


**Figure S7.** Principal component analysis (PCA) for Gs-coupled systems with separated protein components. Magnitudes of the first ten eigenvalues are plotted.

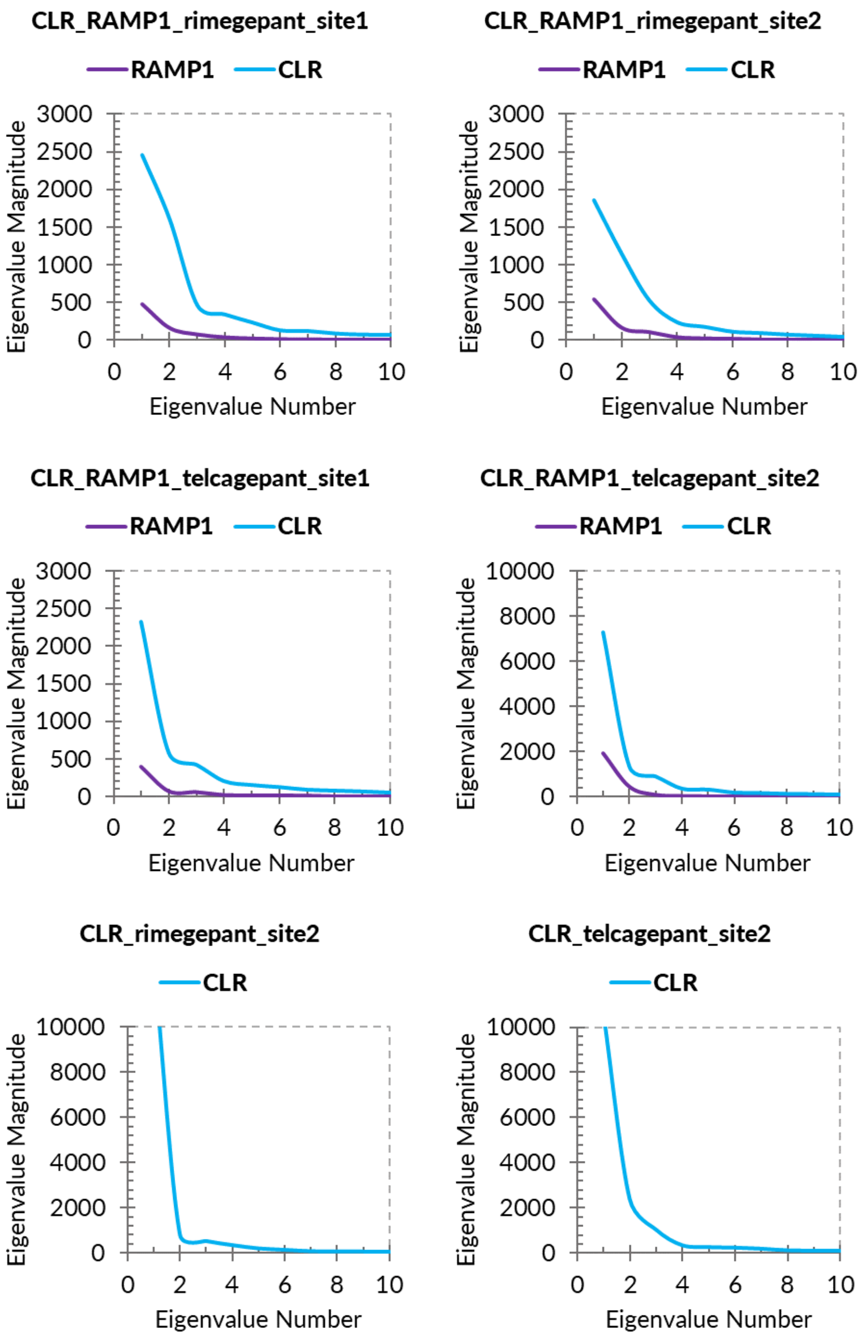


**Figure S8.** Principal component analysis (PCA) for Gs-uncoupled systems with separated protein components. Magnitudes of the first ten eigenvalues are plotted.

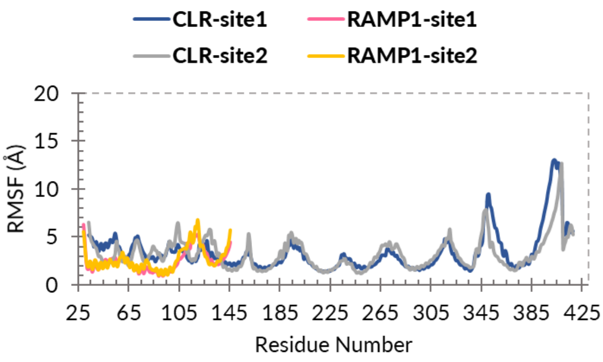


*RMSF_CLR_RAMP1_rimegepant*

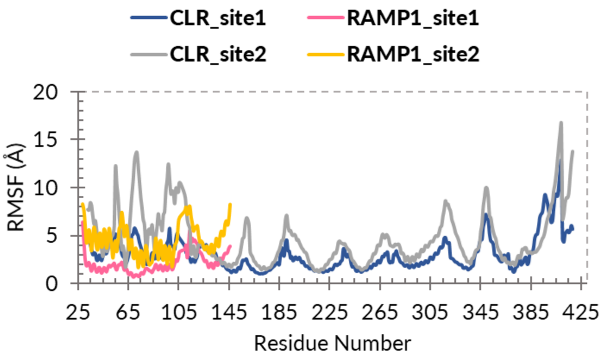


*RMSF_CLR_RAMP1_telcagepant*

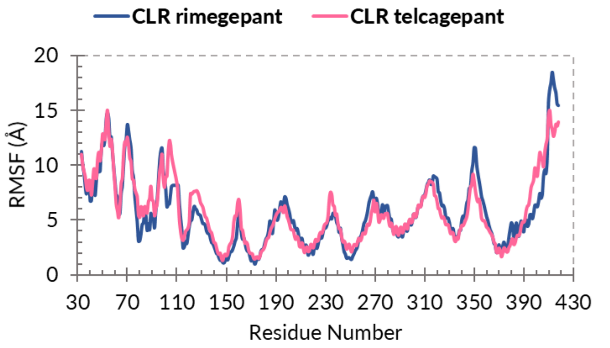


*RMSF_CLR_rimegepant_telcagepant – site2*

**Figure S9.** The Root Mean Square Fluctuation (RMSF) plots of Gs-uncoupled systems calculated using concatenated MD simulation trajectories for each system. RMSF is useful for characterizing local changes along the protein chain. Peaks indicate areas of the protein that fluctuate the most during the simulation. Typically, you will observe that the tails (N- and C-terminal) fluctuate more than any other part of the protein. Secondary structure elements like alpha helices and beta strands are usually more rigid than the unstructured part of the protein, and thus fluctuate less than the loop regions. RMSF value trends of CLR and RAMP1 structure are plotted together (top and middle) for the comparison of the effects of bound gepant molecules at two binding pockets.

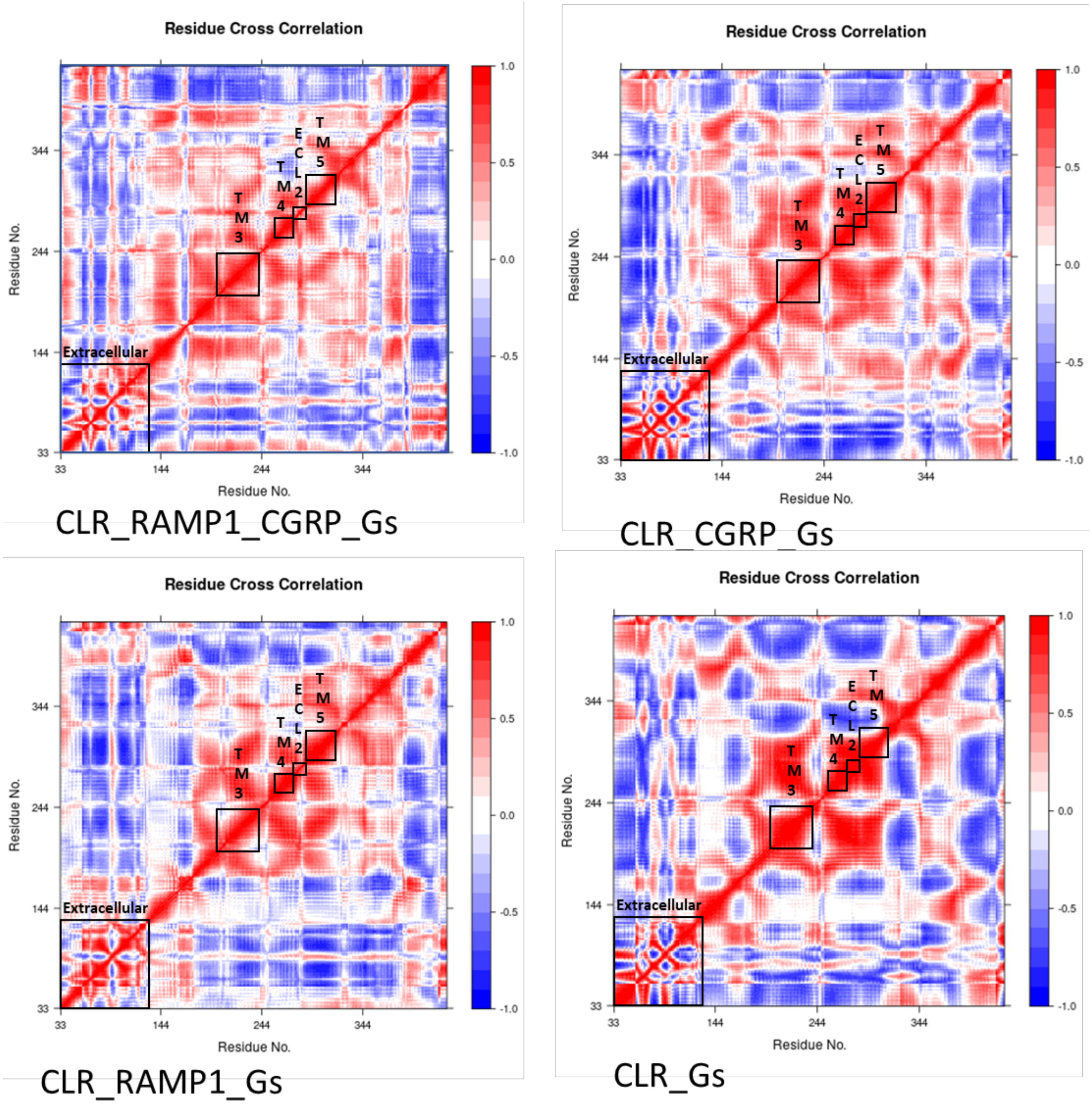


**Figure S10.** Residue cross-correlation analyses of CLR structure in Gs-coupled systems. The extent to which the atomic fluctuations/displacements of a system are correlated with one another can be assessed by examining the magnitude of all pairwise cross-correlation coefficients. The color ladder red to blue showing the correlated residues (red), neutral (white) and anticorrelated residues (blue). Black square frames on the maps are showing the locations of extracellular, TMs 3,4,5 and ECL2 domains to ease the visibility.

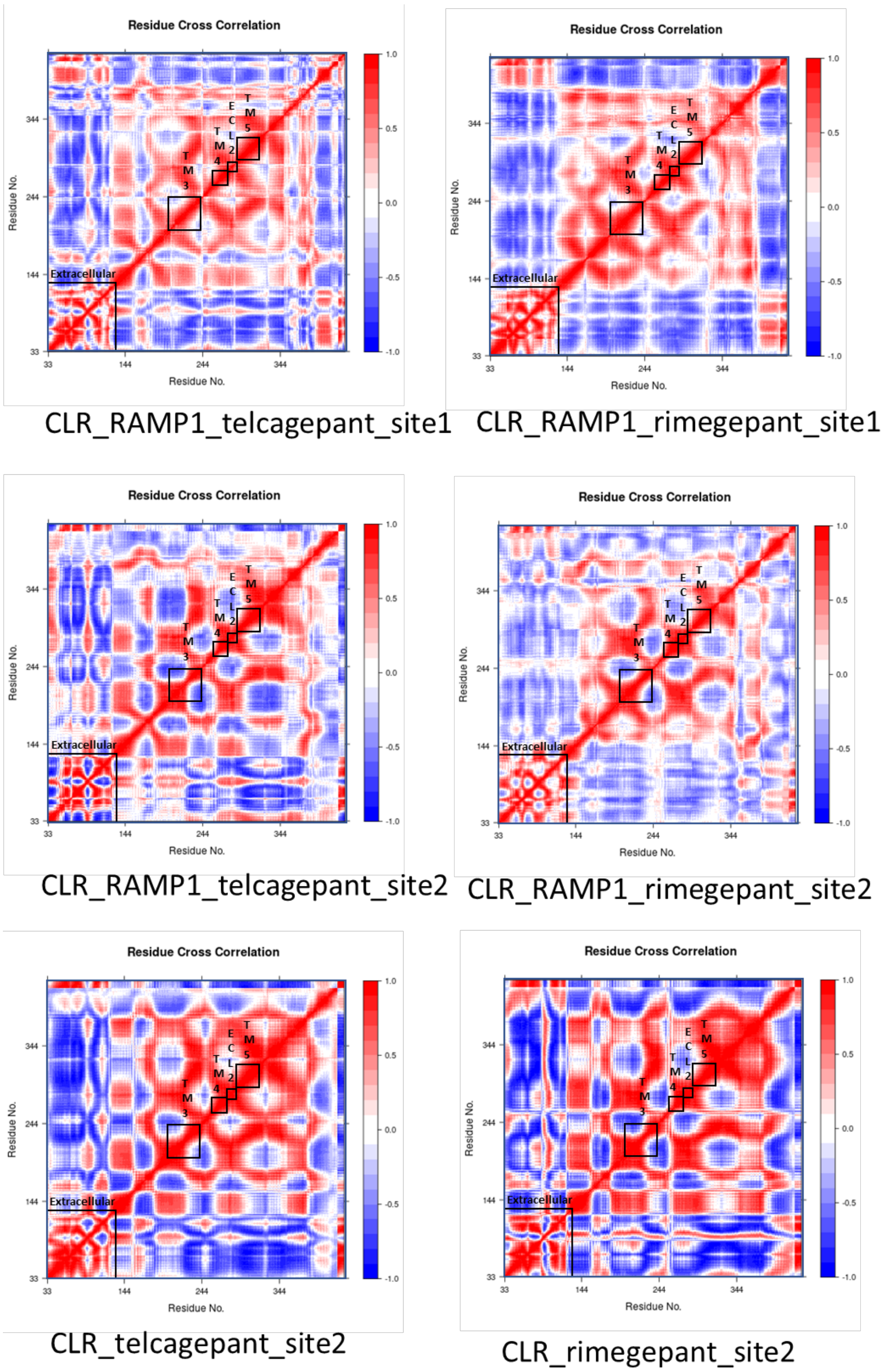


**Figure S11.** Residue cross-correlation analyses of CLR structure in Gs-uncoupled systems. The extent to which the atomic fluctuations/displacements of a system are correlated with one another can be assessed by examining the magnitude of all pairwise cross-correlation coefficients. The color ladder red to blue showing the correlated residues (red), neutral (white) and anticorrelated residues (blue). Black square frames on the maps are showing the locations of extracellular, TMs 3,4,5 and ECL2 domains to ease the visibility.

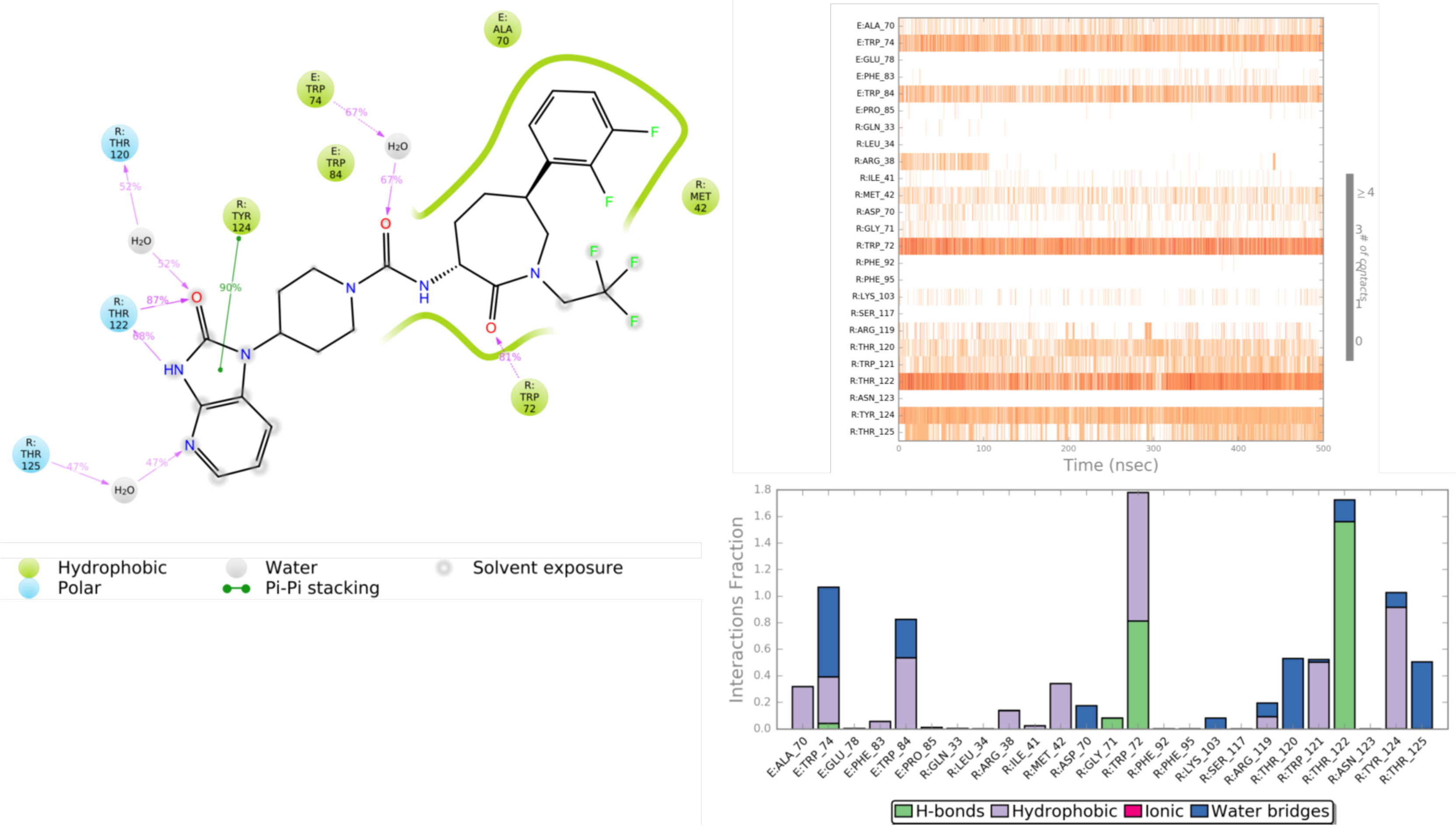


**Figure S12.** The schematic of detailed ligand atom interactions of telcagepant molecule bound at CLR binding ***site 1*** (with RAMP1). Interactions that occur more than 30.0% of the simulation time in the selected trajectory (left). The timeline representation of the interactions and contacts (top right). This panel shows which residues interact with the ligand in each trajectory frame. Some residues make more than one specific contact with the ligand, which is represented by a darker shade of orange, according to the scale to the right of the plot. Protein-Ligand interactions histogram (bottom right).These interactions can be categorized by type and summarized, as shown in the plot above. Protein-ligand interactions are categorized into four types: Hydrogen Bonds (green), Hydrophobic (purple), Ionic (pink) and Water Bridges (blue). The stacked bar charts are normalized over the course of the trajectory: for example, a value of 0.7 suggests that 70% of the simulation time the specific interaction is maintained. Values over 1.0 are possible as some protein residue may make multiple contacts of same subtype with the ligand.

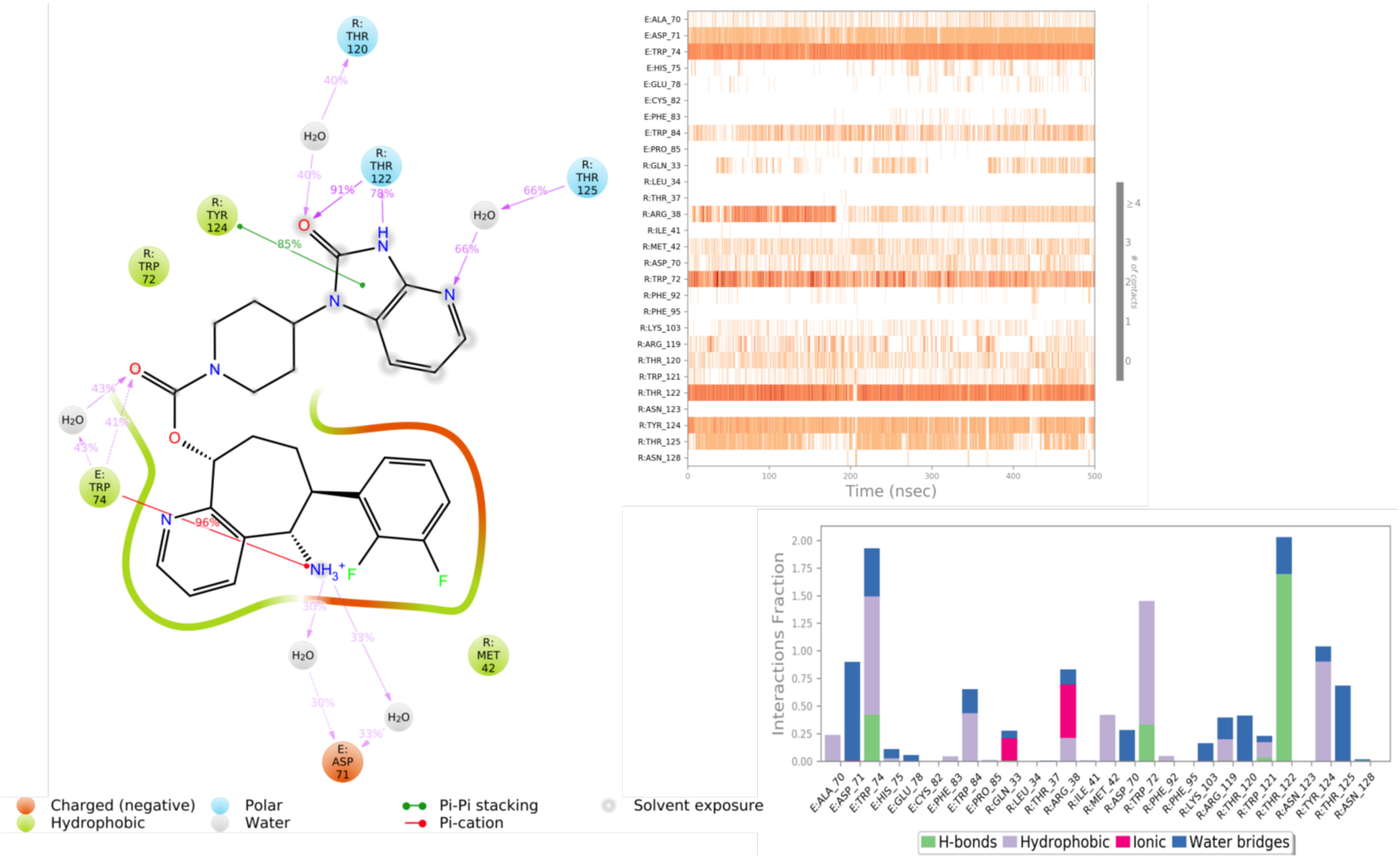


**Figure S13.** The schematic of detailed ligand atom interactions of rimegepant molecule bound at binding ***site 1*** (with RAMP1). Interactions that occur more than 30.0% of the simulation time in the selected trajectory (left). The timeline representation of the interactions and contacts (top right). This panel shows which residues interact with the ligand in each trajectory frame. Some residues make more than one specific contact with the ligand, which is represented by a darker shade of orange, according to the scale to the right of the plot. Protein-Ligand interactions histogram (bottom right).These interactions can be categorized by type and summarized, as shown in the plot above. Protein-ligand interactions are categorized into four types: Hydrogen Bonds (green), Hydrophobic (purple), Ionic (pink) and Water Bridges (blue). The stacked bar charts are normalized over the course of the trajectory: for example, a value of 0.7 suggests that 70% of the simulation time the specific interaction is maintained. Values over 1.0 are possible as some protein residue may make multiple contacts of same subtype with the ligand.

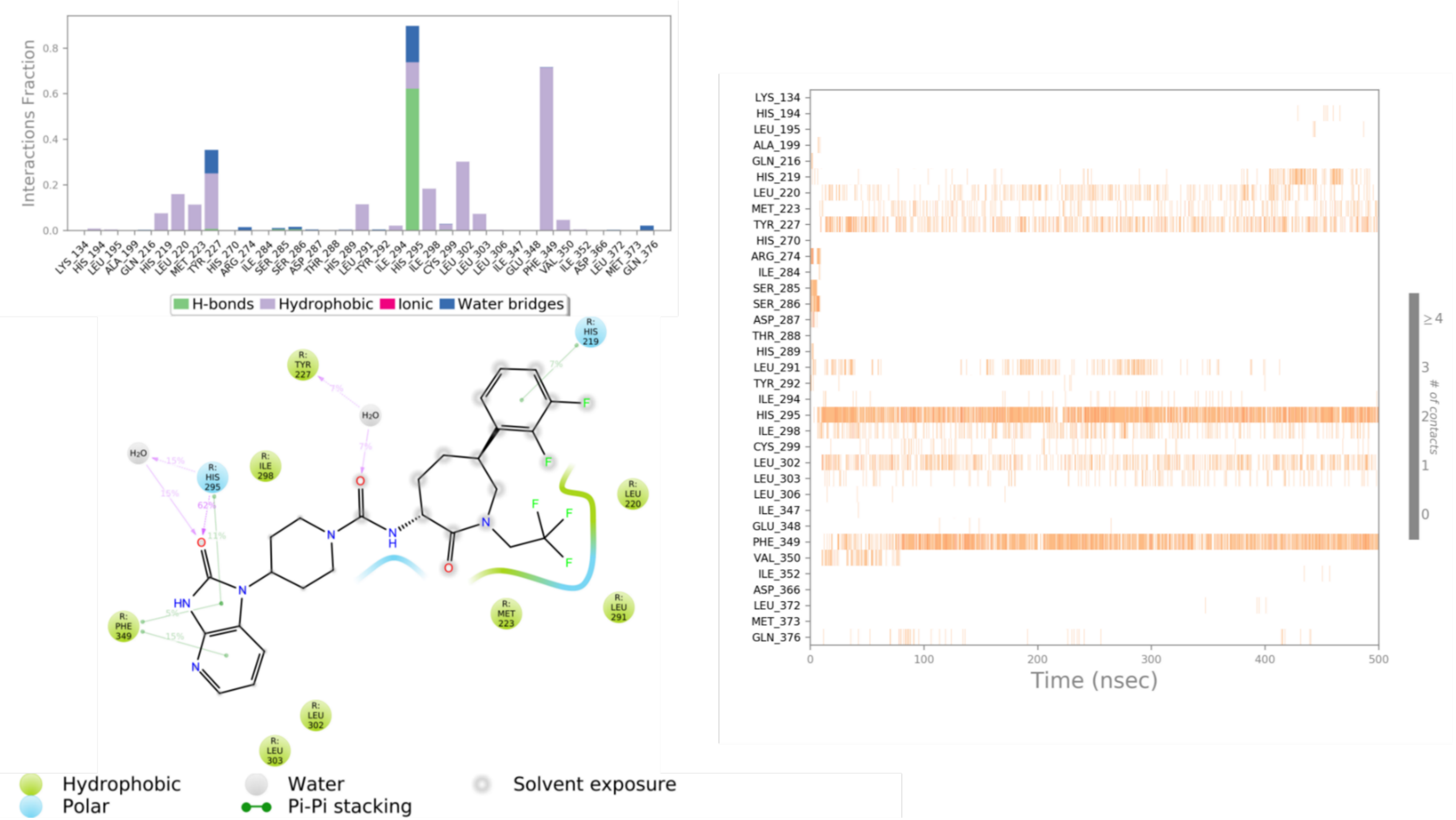


**Figure S14.** The schematic of detailed ligand atom interactions of telcagepant molecule bound at binding ***site 2*** (the first simulation, with RAMP1). Interactions that occur more than 5.0% of the simulation time in the selected trajectory (bottom left). The timeline representation of the interactions and contacts (right). This panel shows which residues interact with the ligand in each trajectory frame. Some residues make more than one specific contact with the ligand, which is represented by a darker shade of orange, according to the scale to the right of the plot. Protein-Ligand interactions histogram (top left).These interactions can be categorized by type and summarized, as shown in the plot above. Protein-ligand interactions are categorized into four types: Hydrogen Bonds (green), Hydrophobic (purple), Ionic (pink) and Water Bridges (blue). The stacked bar charts are normalized over the course of the trajectory: for example, a value of 0.7 suggests that 70% of the simulation time the specific interaction is maintained. Values over 1.0 are possible as some protein residue may make multiple contacts of same subtype with the ligand.

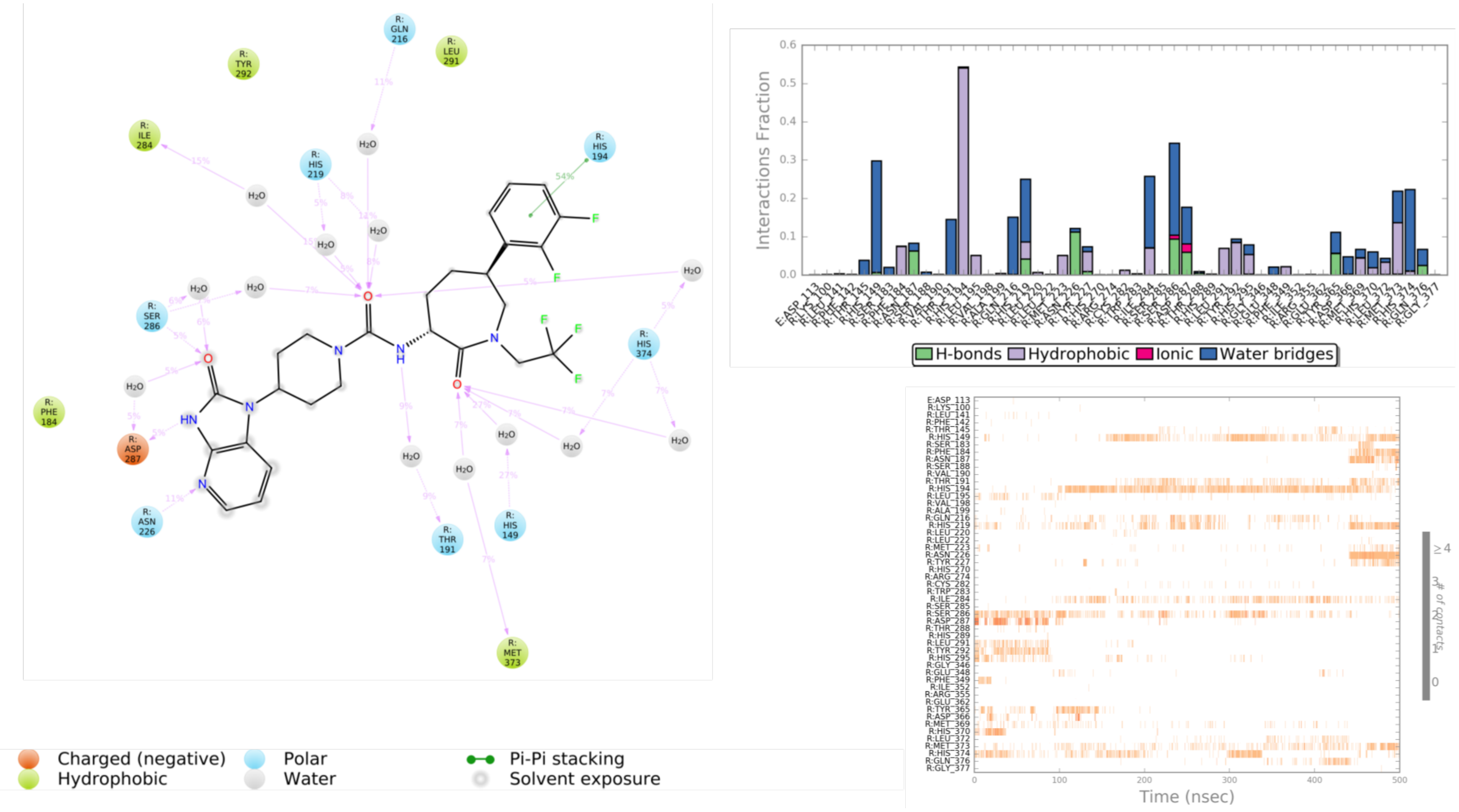


**Figure S14 (cont.).** The schematic of detailed ligand atom interactions of telcagepant molecule bound at binding ***site 2*** (the second simulation, with RAMP1).

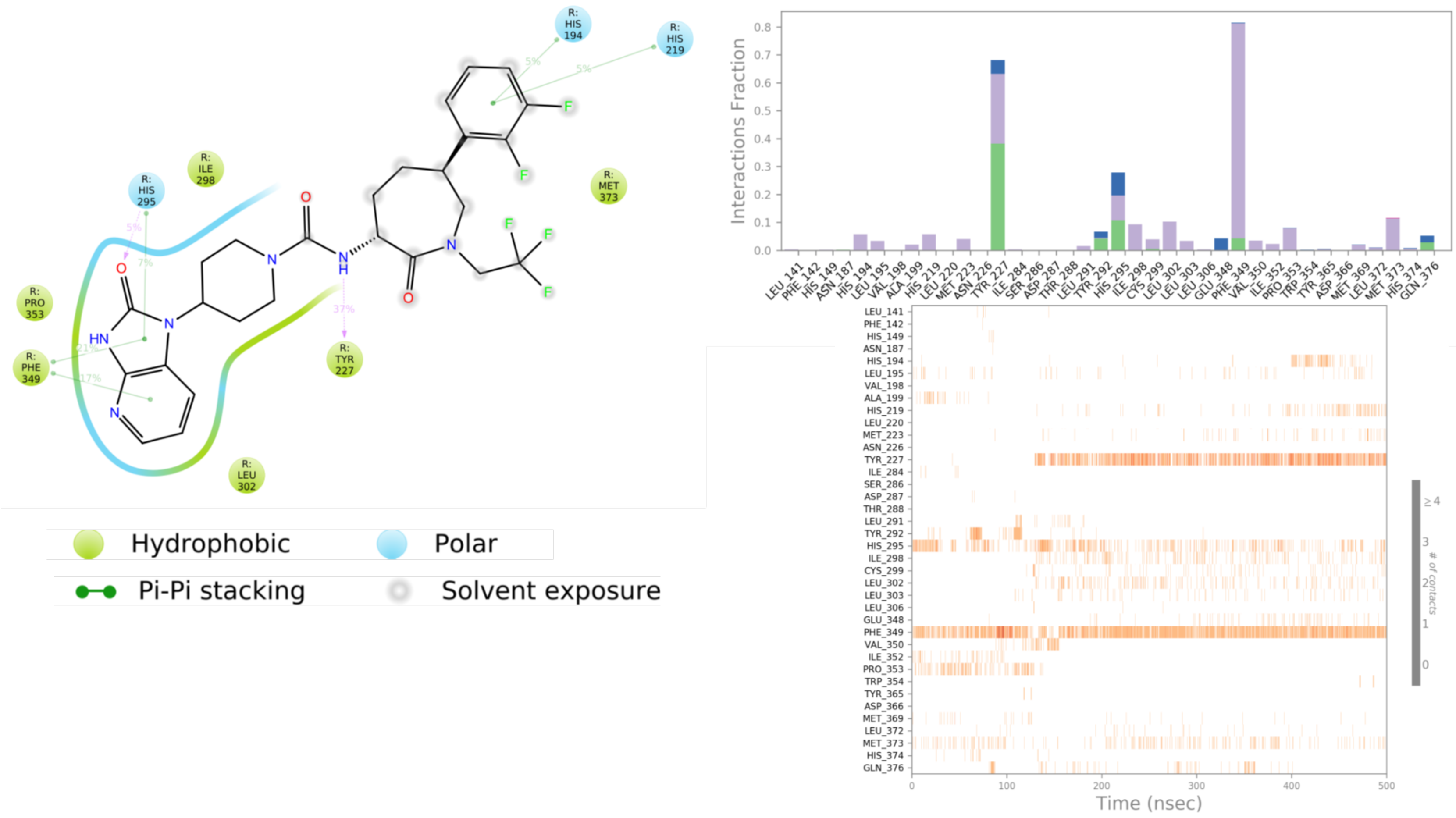


**Figure S14 (cont.).** The schematic of detailed ligand atom interactions of telcagepant molecule bound at binding ***site 2*** (the third simulation, with RAMP1).

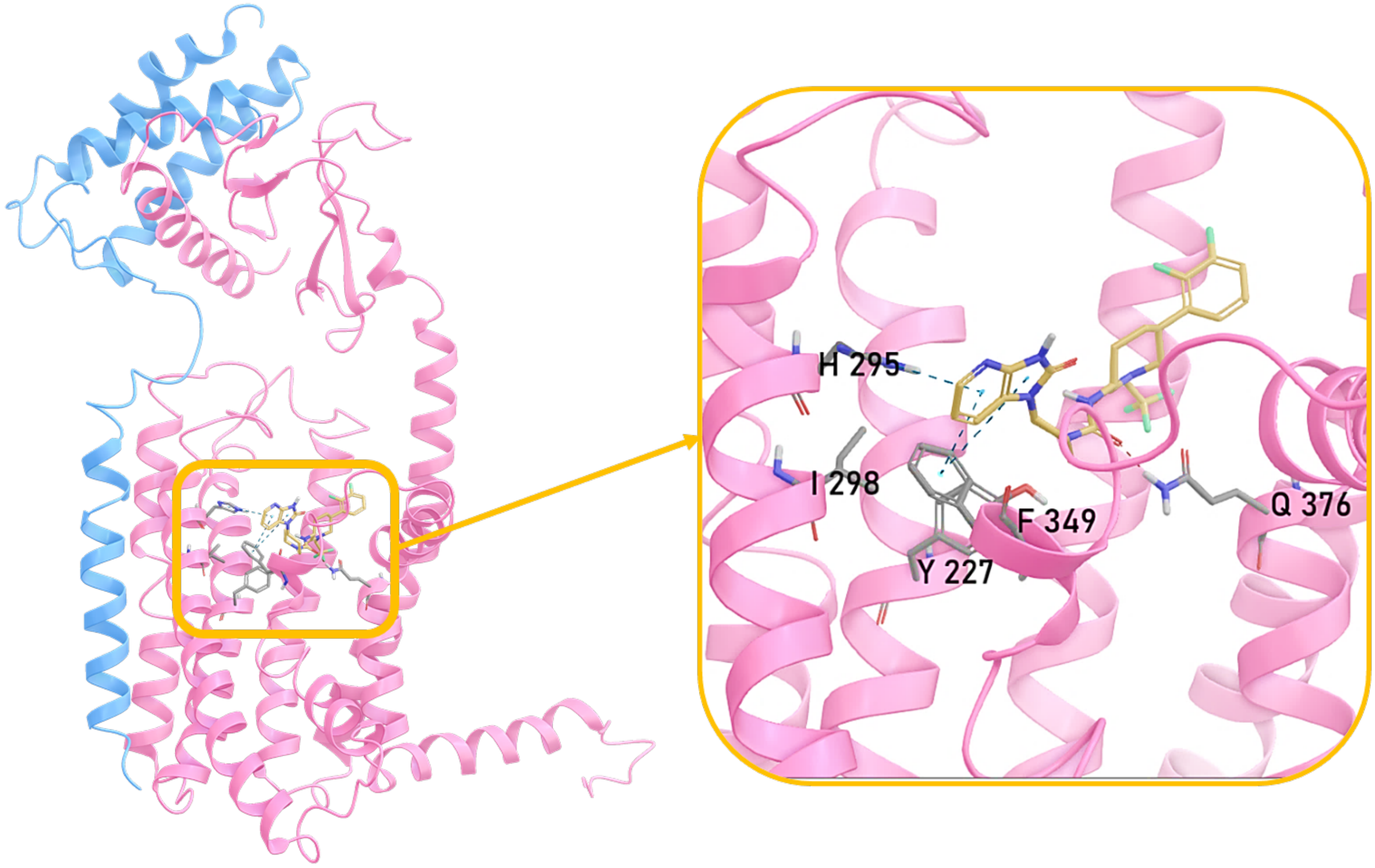


**Figure S14 (cont.).** Rep structure extracted from concatenated trajectories of telcagepant molecule bound at binding ***site 2.*** Protein structures are shown in ribbon representation. RAMP1 structures are shown in blue ribbons, CLR structures are shown in pink colors. In zoomed views, ligand and interacting residues are shown in stick representation. Dashed lines show the H-bonds, salt bridges and π-π stacking interactions. We extracted the representative structures from the concatenated MD simulation trajectories by choosing the one with the closest RMSD to the average structure.

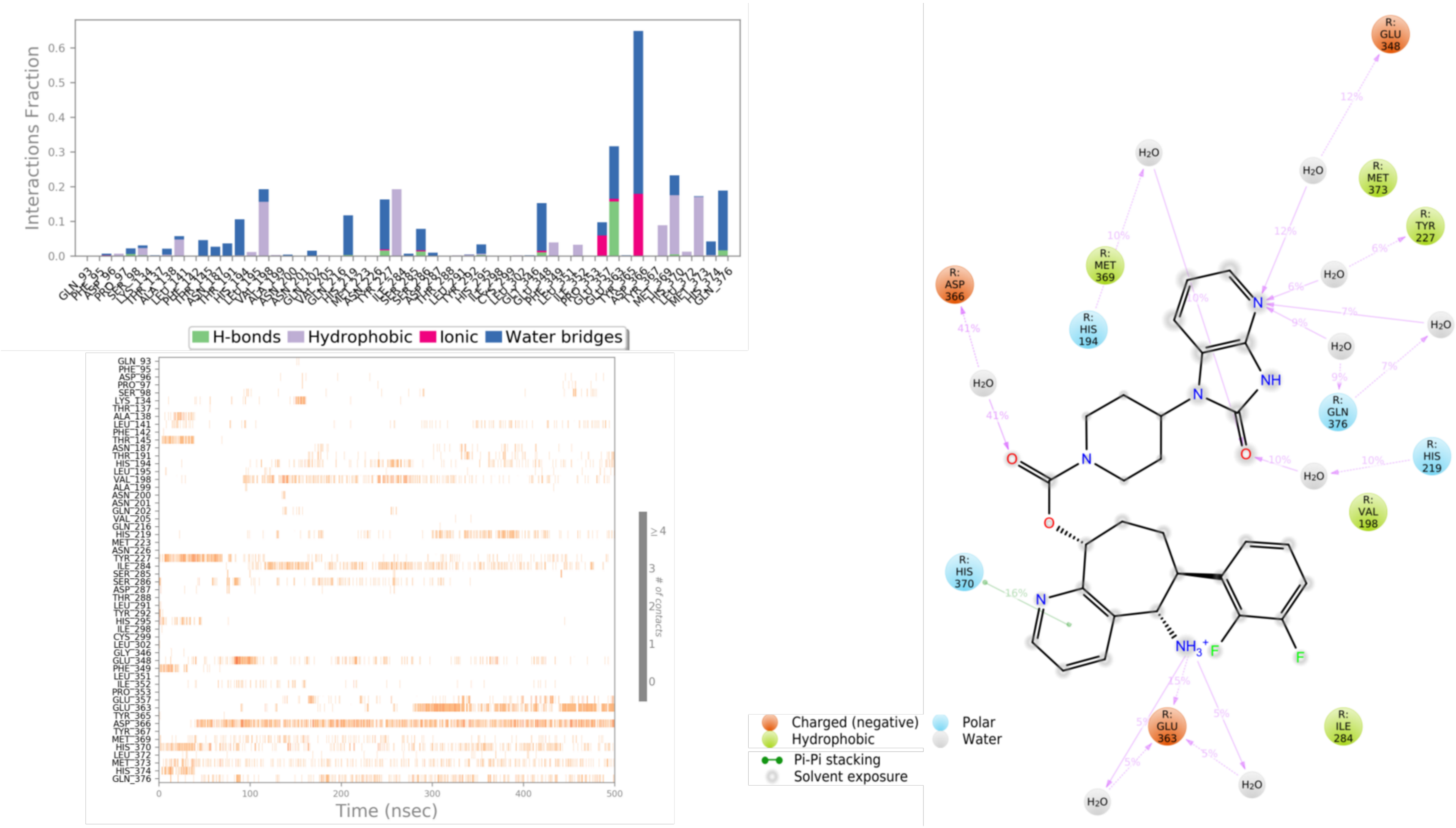

**Figure S15.** The schematic of detailed ligand atom interactions of rimegepant molecule bound at binding ***site 2*** (the first simulation, with RAMP1). Interactions that occur more than 5.0% of the simulation time in the selected trajectory (right). The timeline representation of the interactions and contacts (bottom left). This panel shows which residues interact with the ligand in each trajectory frame. Some residues make more than one specific contact with the ligand, which is represented by a darker shade of orange, according to the scale to the right of the plot. Protein-Ligand interactions histogram (top left).These interactions can be categorized by type and summarized, as shown in the plot above. Protein-ligand interactions are categorized into four types: Hydrogen Bonds (green), Hydrophobic (purple), Ionic (pink) and Water Bridges (blue). The stacked bar charts are normalized over the course of the trajectory: for example, a value of 0.7 suggests that 70% of the simulation time the specific interaction is maintained. Values over 1.0 are possible as some protein residue may make multiple contacts of same subtype with the ligand.

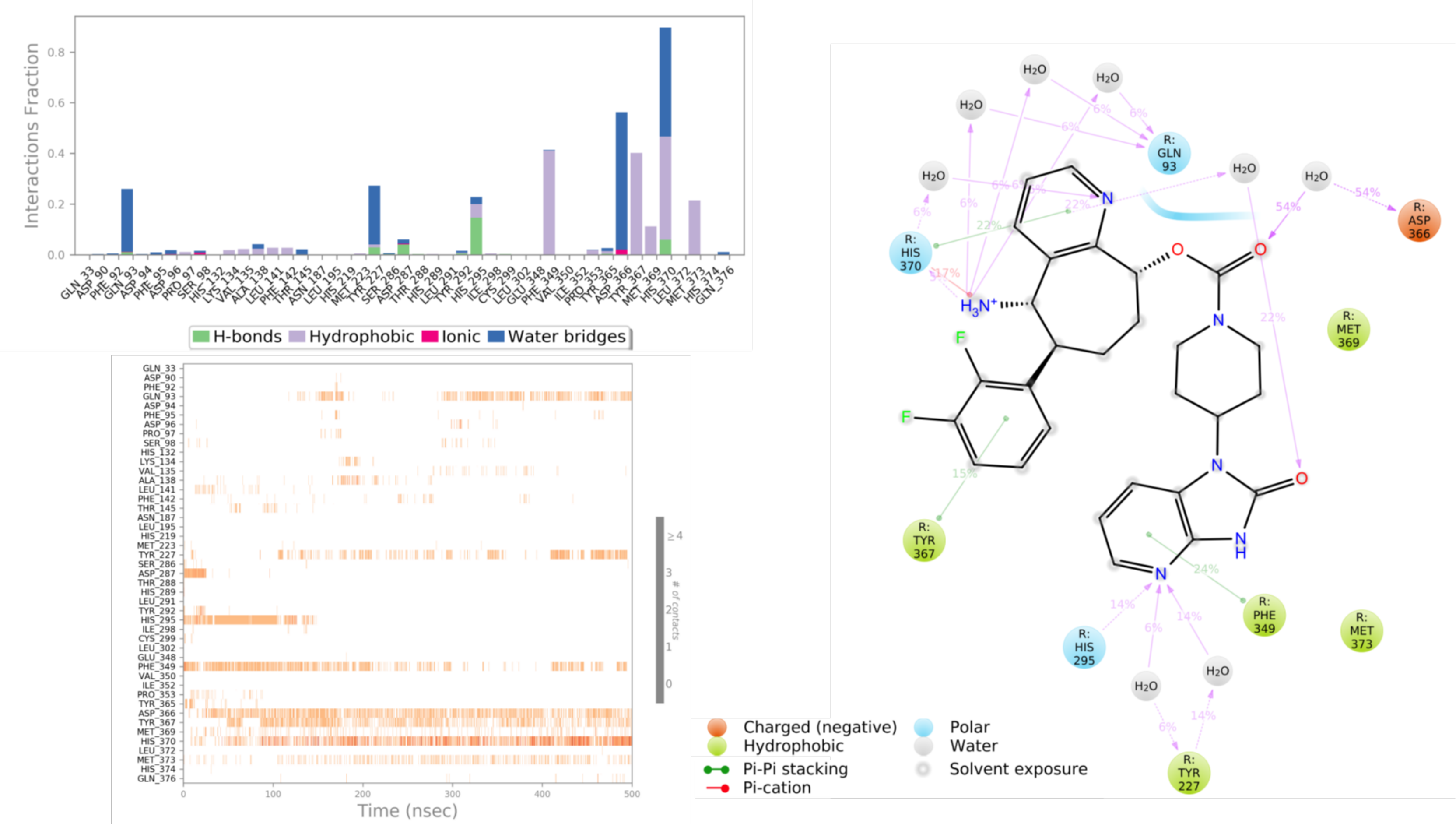


**Figure S15 (cont.).** The schematic of detailed ligand atom interactions of rimegepant molecule bound at binding ***site 2*** (the second simulation, with RAMP1).

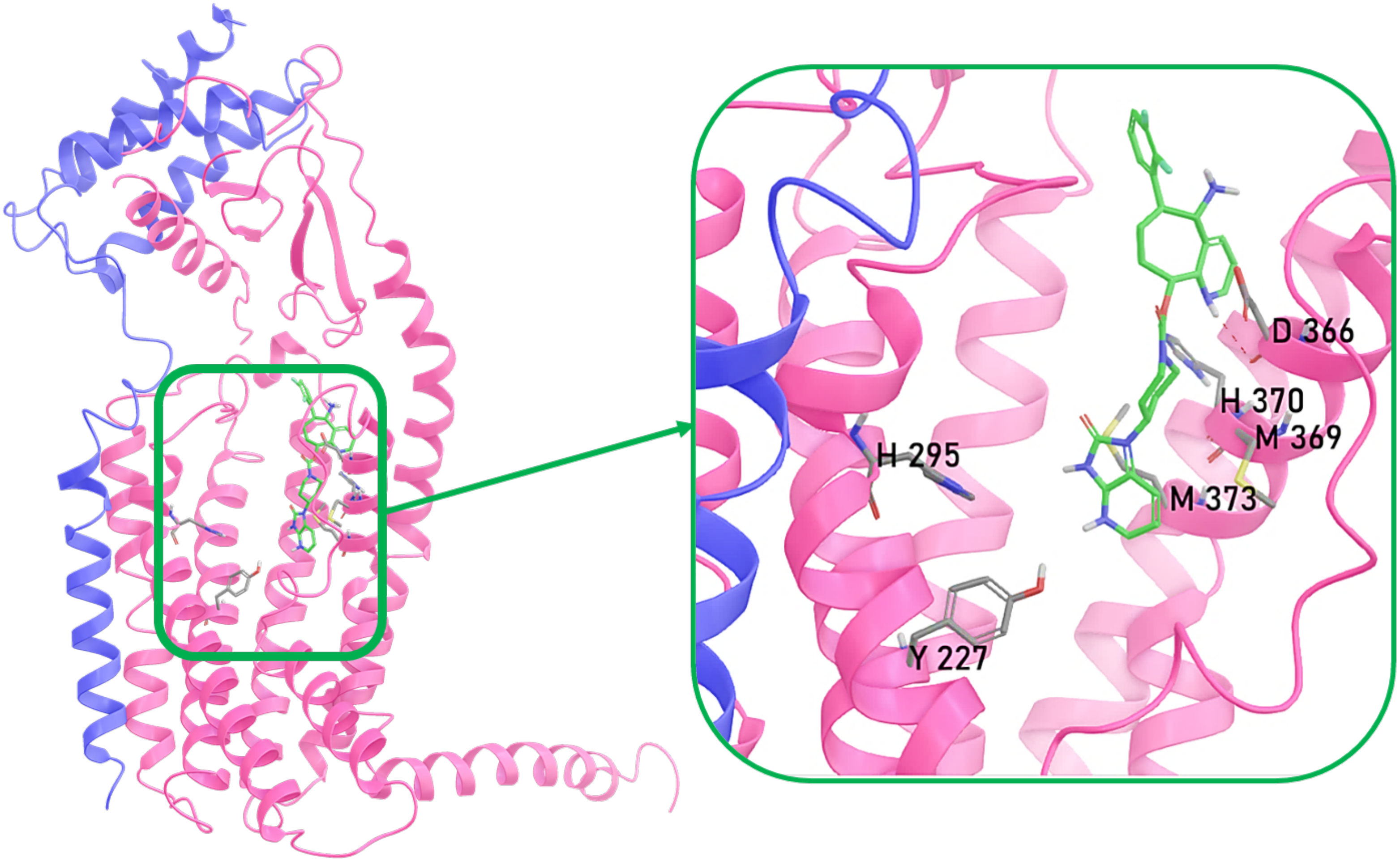


**Figure S15 (cont.).** Rep structure extracted from concatenated trajectories of rimegepant molecule bound at binding ***site 2.*** Protein structures are shown in ribbon representation. RAMP1 structures are shown in blue ribbons, CLR structures are shown in pink colors. In zoomed views, ligand and interacting residues are shown in stick representation. Dashed lines show the H-bonds, salt bridges and π-π stacking interactions. We extracted the representative structures from the concatenated MD simulation trajectories by choosing the one with the closest RMSD to the average structure.

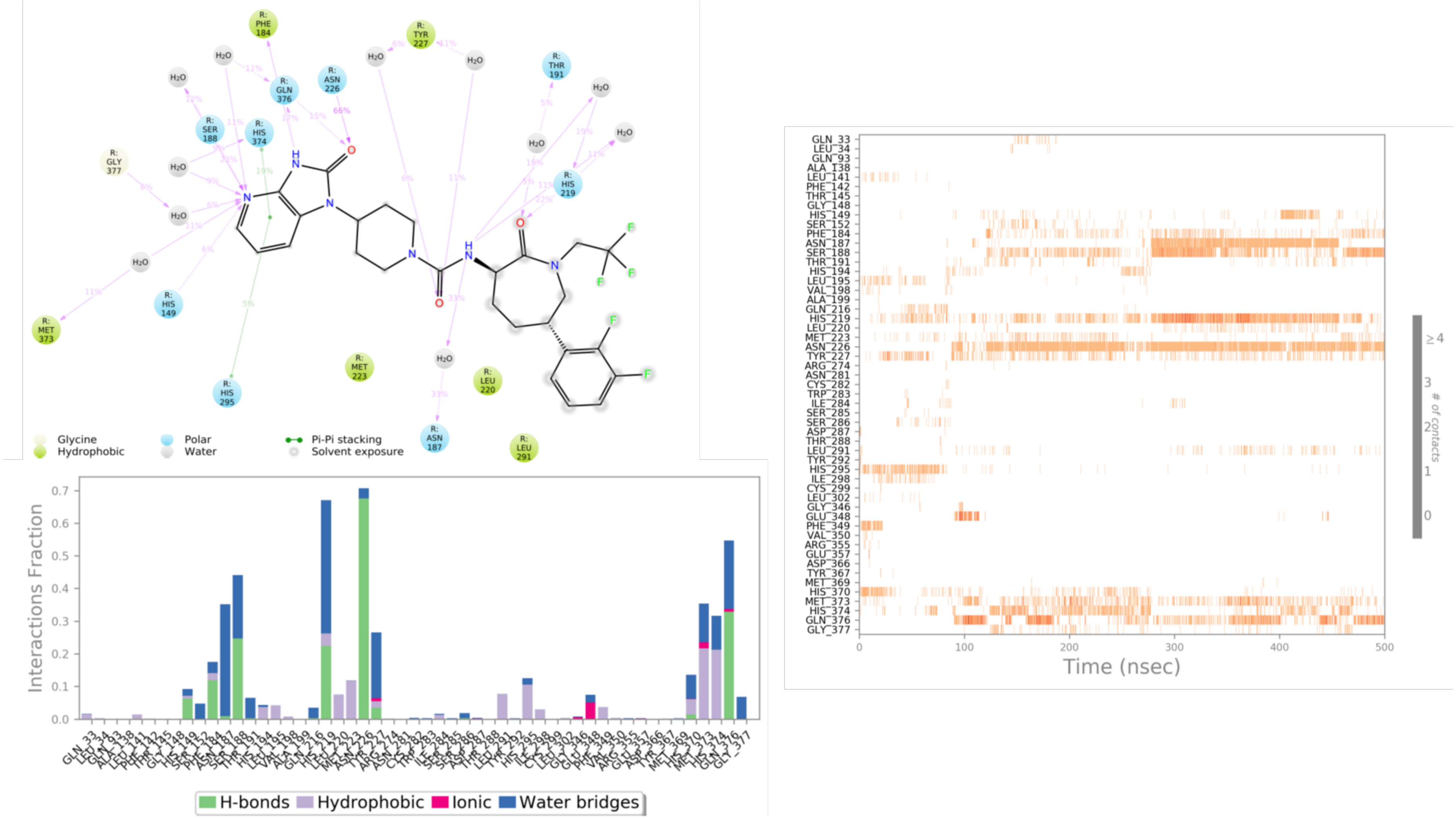


**Figure S16.** The schematic of detailed ligand atom interactions of telcagepant molecule bound at CLR binding ***site 2*** (the first simulation, without RAMP1). Interactions that occur more than 5.0% of the simulation time in the selected trajectory (top left). The timeline representation of the interactions and contacts (right). This panel shows which residues interact with the ligand in each trajectory frame. Some residues make more than one specific contact with the ligand, which is represented by a darker shade of orange, according to the scale to the right of the plot. Protein-Ligand interactions histogram (bottom left).These interactions can be categorized by type and summarized, as shown in the plot above. Protein-ligand interactions are categorized into four types: Hydrogen Bonds (green), Hydrophobic (purple), Ionic (pink) and Water Bridges (blue). The stacked bar charts are normalized over the course of the trajectory: for example, a value of 0.7 suggests that 70% of the simulation time the specific interaction is maintained. Values over 1.0 are possible as some protein residue may make multiple contacts of same subtype with the ligand.

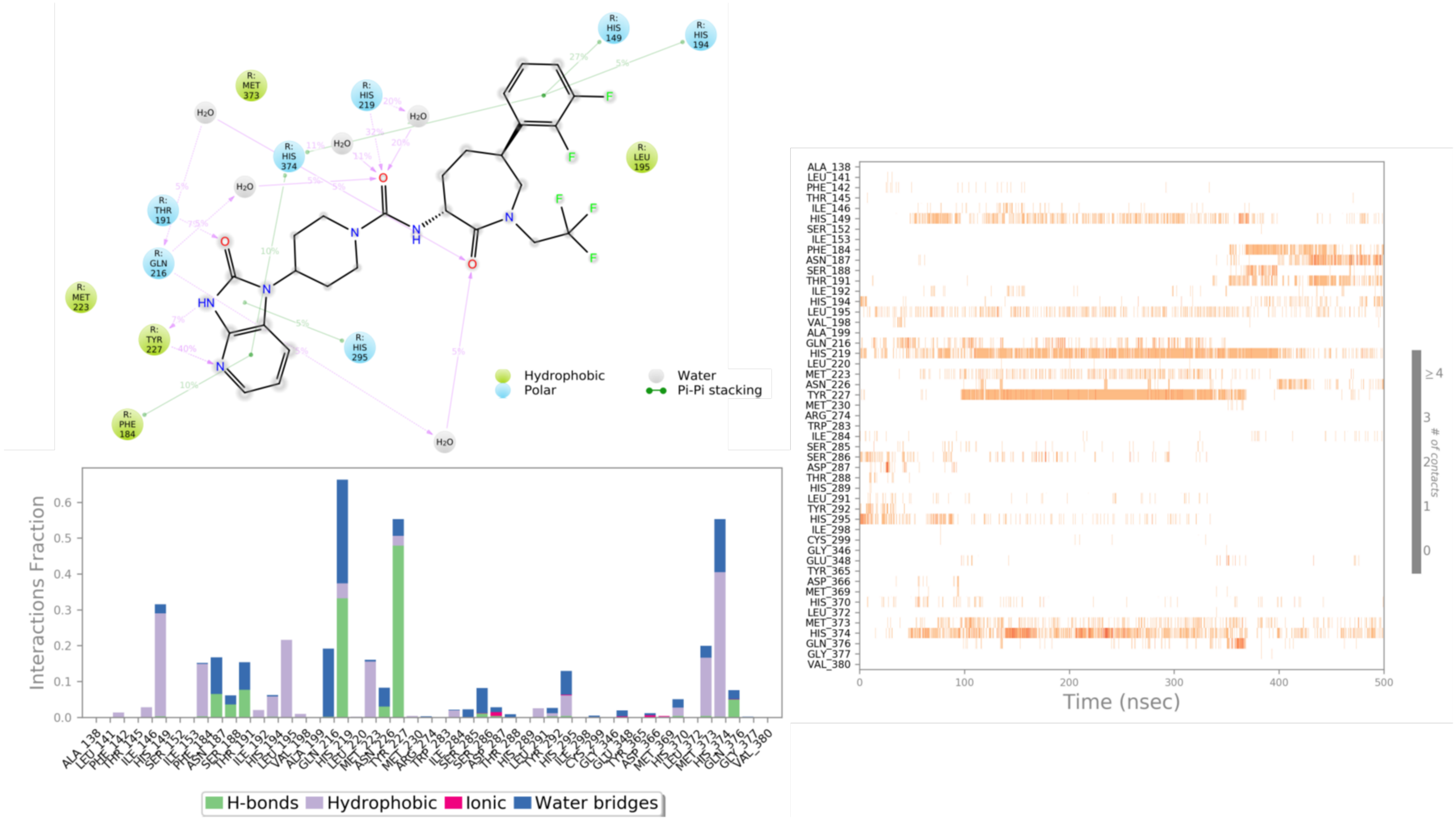

**Figure S16 (cont.).** The schematic of detailed ligand atom interactions of telcagepant molecule bound at CLR binding ***site 2*** (the second simulation, without RAMP1).

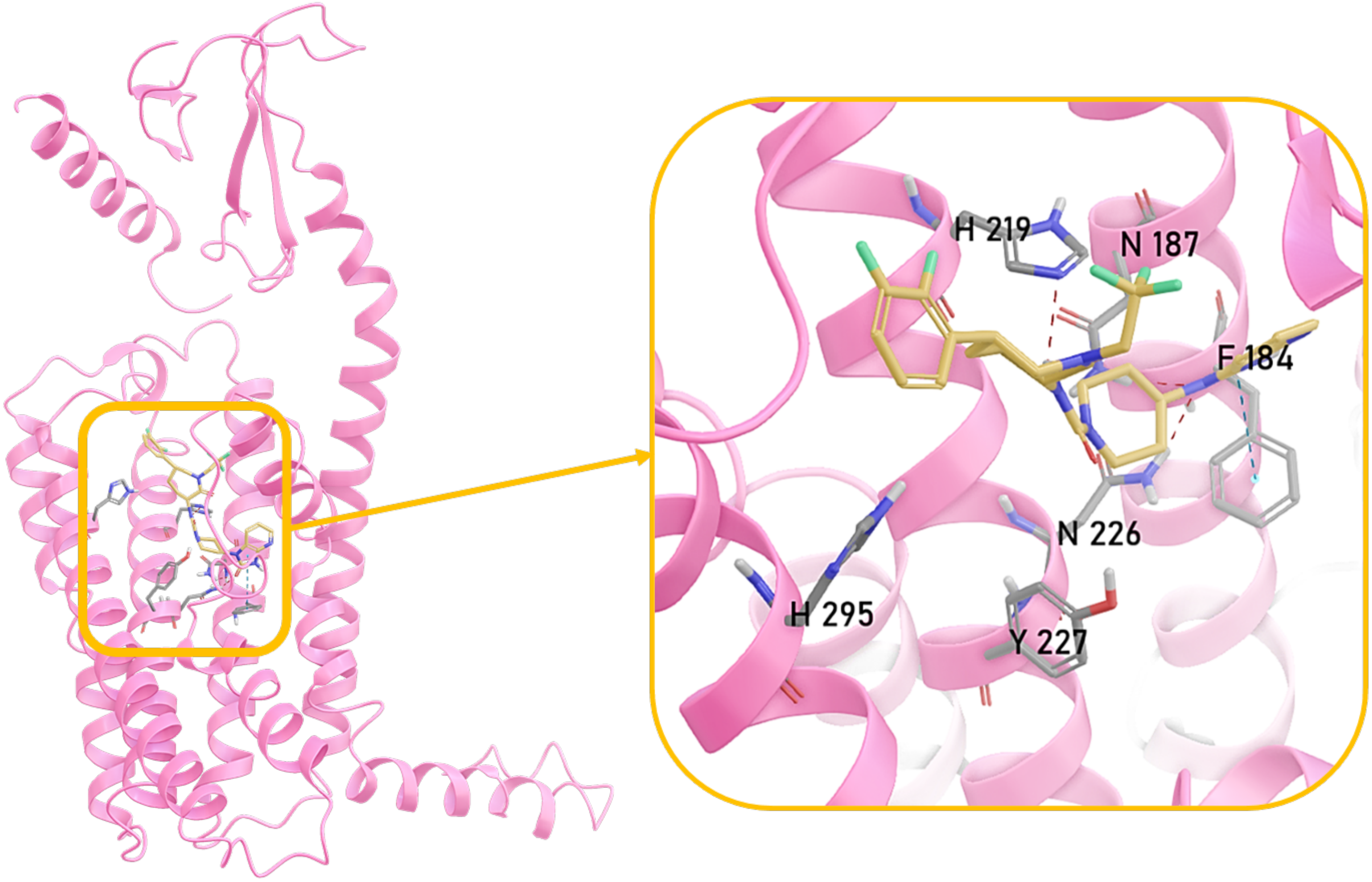


**Figure S16 (cont.).** Rep structure extracted from concatenated trajectories of telcagepant molecule bound at binding ***site 2.*** CLR structure is shown in pink ribbon representation. In zoomed views, ligand molecule and interacting residues are shown in stick representation. Dashed lines show the H-bonds, salt bridges and π-π stacking interactions. We extracted the representative structures from the concatenated MD simulation trajectories by choosing the one with the closest RMSD to the average structure.

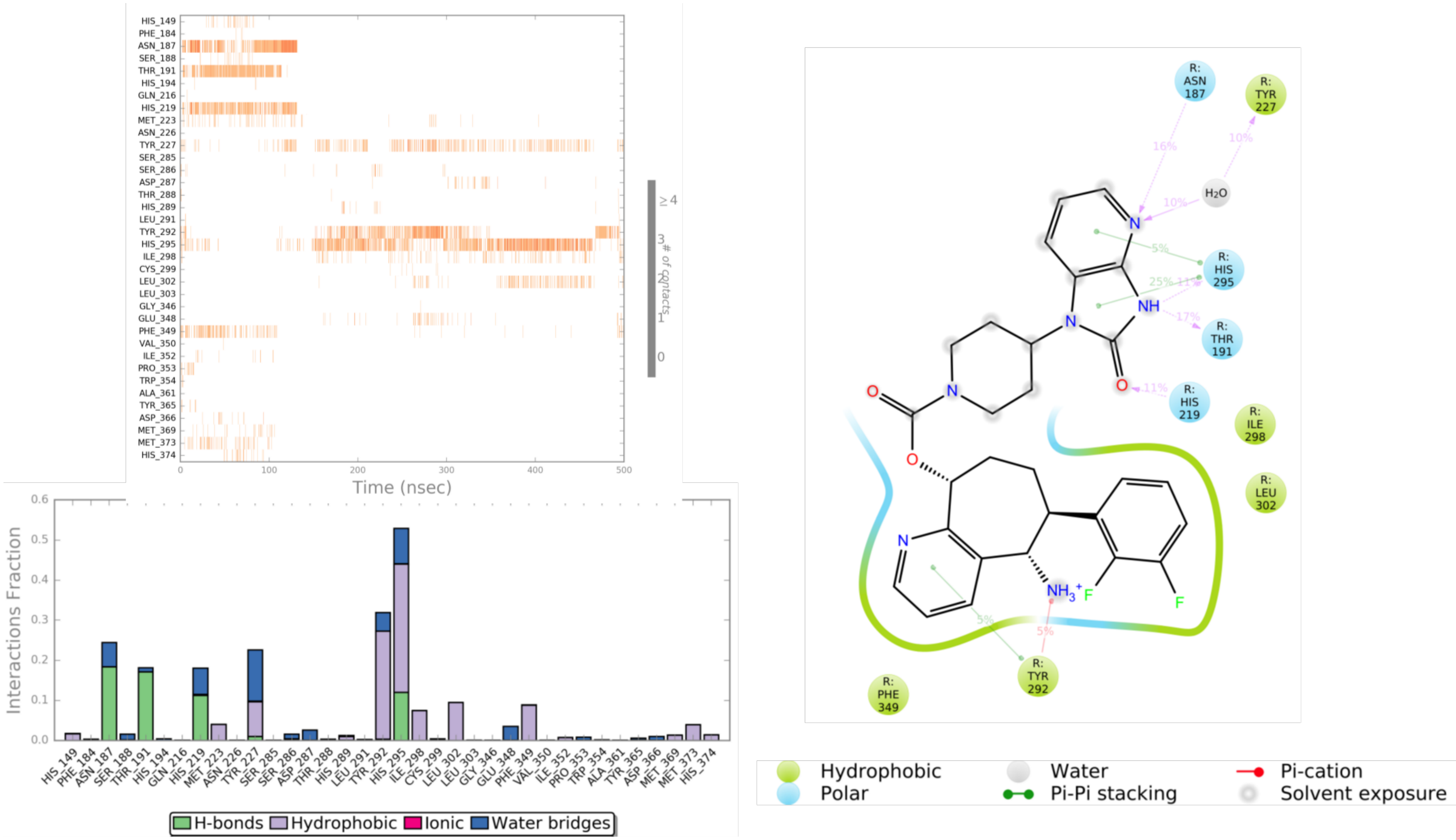


**Figure S17.** The schematic of detailed ligand atom interactions of rimegepant molecule bound at CLR binding ***site 2*** (the first simulation, without RAMP1). Interactions that occur more than 5.0% of the simulation time in the selected trajectory (right). The timeline representation of the interactions and contacts (top left). This panel shows which residues interact with the ligand in each trajectory frame. Some residues make more than one specific contact with the ligand, which is represented by a darker shade of orange, according to the scale to the right of the plot. Protein-Ligand interactions histogram (bottom left).These interactions can be categorized by type and summarized, as shown in the plot above. Protein-ligand interactions are categorized into four types: Hydrogen Bonds (green), Hydrophobic (purple), Ionic (pink) and Water Bridges (blue). The stacked bar charts are normalized over the course of the trajectory: for example, a value of 0.7 suggests that 70% of the simulation time the specific interaction is maintained. Values over 1.0 are possible as some protein residue may make multiple contacts of same subtype with the ligand.

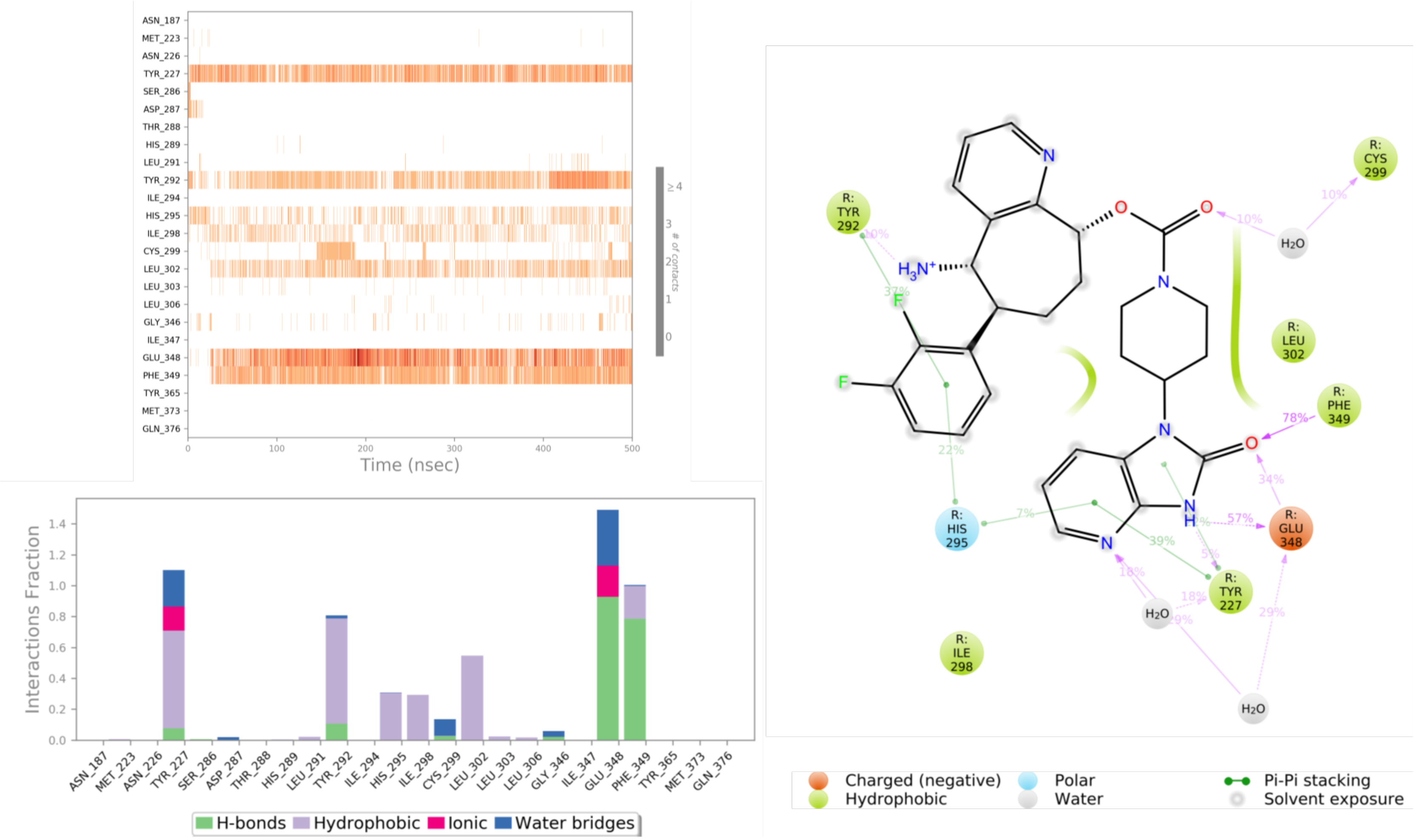


**Figure S17 (cont.).** The schematic of detailed ligand atom interactions of rimegepant molecule bound at CLR binding ***site 2*** (the second simulation, without RAMP1).

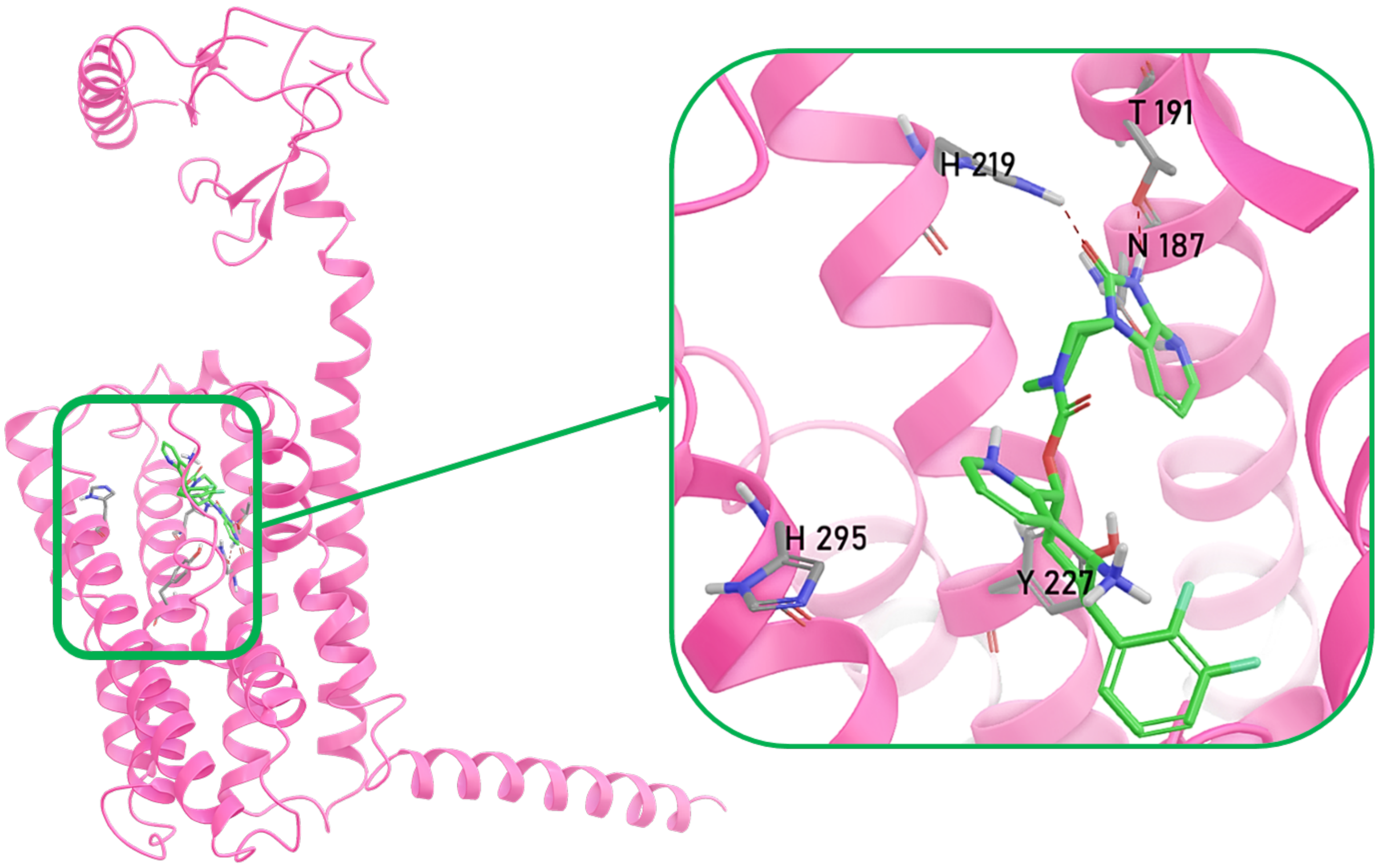


**Figure S17 (cont).** Rep structure extracted from concatenated trajectories of rimegepant molecule bound at binding ***site 2.*** CLR structure is shown in pink ribbon representation. In zoomed views, ligand molecule and interacting residues are shown in stick representation. Dashed lines show the H-bonds, salt bridges and π-π stacking interactions. We extracted the representative structures from the concatenated MD simulation trajectories by choosing the one with the closest RMSD to the average structure.

**Supplementary Tables**

**Table S1.** Average MM/GBSA dG – NS values for all MD replicas

| System | Average MM/GBSA dG – NS (kcal/mol) | | |
|---|---|---|---|
| | replica1 | replica2 | replica3 |
| CLR_RAMP1_rimegepant_site 1 | -99.0 ± 6.9 | -101.1 ± 6.8 | n/a |
| CLR_RAMP1_telcagepant_site 1 | -106.8 ± 6.3 | -104.5 ± 7.9 | n/a |
| CLR_RAMP1_rimegepant_site 2 | -48.5 ± 7.0 | -51.6 ± 6.1 | n/a |
| CLR_RAMP1_telcagepant_site 2 | -68.0 ± 7.8 | -56.5 ± 12.7 | -55.5 ± 7.7 |
| CLR_rimegepant_site2 | -40.1 ± 8.6 | -57.7 ± 6.4 | n/a |
| CLR_telcagepant_site2 | -63.6 ± 12.1 | -55.6 ± 9.2 | n/a |

**Table S2.** The frequency table of nonpolar interactions (VdW and pi – pi stacking) between CGRP neuropeptide and CLR with or without RAMP1 structure. The interactions occurred less than 30% of total concatenated trajectories are not shown (n/a).

| CLR Domain | Residue Pair | without RAMP1 | with RAMP1 |
|---|---|---|---|
| ECD | CLR D94 - CGRP P29 | 36.3% | 73.7% |
| ECD | CLR D94 - CGRP T30 | 39.5% | 96.1% |
| ECD | CLR D96 - CGRP R18 | n/a | 33.6% |
| ECD | CLR F92 - CGRP T30 | 40.1% | 52.1% |
| ECD | CLR F95 - CGRP R18 | n/a | 31.7% |
| ECD | CLR F95 - CGRP T30 | 34.3% | 38.3% |
| ECD | CLR G71 - CGRP F37 | n/a | 38.6% |
| ECD | CLR H114 - CGRP S34 | n/a | 51.1% |
| ECD | CLR H132 - CGRP V22 | 40.8% | n/a |
| ECD | CLR N128 - CGRP T30 | 38.5% | 38.2% |
| ECD | CLR N128 - CGRP V32 | n/a | 43.9% |
| ECD | CLR P97 - CGRP R18 | n/a | 62.7% |
| ECD | CLR Q93 - CGRP R18 | 42.4% | 84.5% |
| ECD | CLR Q93 - CGRP S17 | n/a | 31.4% |
| ECD | CLR Q93 - CGRP S19 | 38.5% | 52.5% |
| ECD | CLR R119 - CGRP F37 | 38.0% | 47.1% |
| ECD | CLR S117 - CGRP S34 | n/a | 54.5% |
| ECD | CLR T120 - CGRP F37 | 42.0% | n/a |
| ECD | CLR T122 - CGRP F37 | 44.1% | 69.1% |
| ECD | CLR T125 - CGRP V32 | n/a | 49.0% |
| ECD | CLR V135 - CGRP L15 | 40.5% | n/a |
| ECD | CLR V135 - CGRP S19 | 36.9% | 58.4% |
| ECD | CLR V135 - CGRP V22 | n/a | 36.9% |
| ECD | CLR V135 - CGRP V23 | 40.6% | 37.0% |
| ECD | CLR W121 - CGRP A36 | 37.2% | 44.1% |
| ECD | CLR W121 - CGRP F37 | 39.0% | n/a |
| ECD | CLR W121 - CGRP F37 | n/a | 46.7% |
| ECD | CLR W121 - CGRP G33 | n/a | 64.3% |
| ECD | CLR W121 - CGRP S34 | n/a | 31.6% |
| ECD | CLR W72 - CGRP F37 | n/a | 57.4% |
| ECD | CLR W72 - CGRP T30 | n/a | 38.4% |
| ECD | CLR W72 - CGRP V32 | n/a | 41.2% |
| ECD | CLR Y124 - CGRP F37 | n/a | 50.5% |
| ECD | CLR Y124 - CGRP V32 | n/a | 54.6% |
| ECD-TM1 | CLR A138 - CGRP L12 | 45.6% | n/a |

| | | | |
|---|---|---|---|
| ECD-TM1 | CLR A138 - CGRP L15 | 47.8% | 51.1% |
| ECD-TM1 | CLR A138 - CGRP L16 | 30.8% | n/a |
| ECD-TM1 | CLR L139 - CGRP L16 | 35.0% | n/a |
| ECL1 | CLR A199 - CGRP L16 | 49.9% | 50.8% |
| ECL1 | CLR A199 - CGRP S17 | n/a | 49.1% |
| ECL1 | CLR N200 - CGRP K24 | n/a | 38.4% |
| ECL1 | CLR N200 - CGRP L16 | n/a | 30.1% |
| ECL1 | CLR Q202 - CGRP G20 | 37.7% | 39.4% |
| ECL1 | CLR Q202 - CGRP G21 | n/a | 55.7% |
| ECL1 | CLR Q202 - CGRP K24 | n/a | 50.7% |
| ECL1 | CLR Q202 - CGRP L16 | 51.0% | 51.1% |
| ECL1 | CLR Q202 - CGRP S17 | 53.0% | 67.9% |
| ECL1 | CLR V198 - CGRP A13 | 40.9% | n/a |
| ECL1 | CLR V198 - CGRP S17 | 42.7% | n/a |
| ECL2 | CLR D280 - CGRP R18 | 33.3% | n/a |
| ECL2 | CLR D287 - CGRP A1 | n/a | 39.8% |
| ECL2 | CLR D287 - CGRP R18 | 44.2% | 32.7% |
| ECL2 | CLR I284 - CGRP A13 | 36.1% | 53.4% |
| ECL2 | CLR I284 - CGRP G14 | 36.5% | n/a |
| ECL2 | CLR I284 - CGRP H10 | 69.5% | n/a |
| ECL2 | CLR I284 - CGRP S17 | 43.3% | 52.2% |
| ECL2 | CLR S286 - CGRP A1 | 39.6% | 34.8% |
| ECL2 | CLR S286 - CGRP H10 | 60.2% | 32.8% |
| ECL2 | CLR S286 - CGRP R11 | 46.6% | 49.7% |
| ECL3 | CLR R355 - CGRP D3 | n/a | 45.6% |
| TM1 | CLR F142 - CGRP L16 | 52.8% | 69.5% |
| TM2 | CLR H194 - CGRP A13 | 52.1% | n/a |
| TM2 | CLR H194 - CGRP H10 | 30.0% | n/a |
| TM2 | CLR H194 - CGRP T9 | 35.1% | n/a |
| TM2 | CLR L195 - CGRP A13 | n/a | 41.3% |
| TM2 | CLR L195 - CGRP L16 | 36.8% | 41.0% |
| TM3 | CLR H219 - CGRP H10 | 35.4% | 39.6% |
| TM3 | CLR H219 - CGRP T9 | n/a | 44.7% |
| TM3 | CLR L220 - CGRP H10 | 46.5% | 51.2% |
| TM3 | CLR M223 - CGRP T6 | 36.2% | 36.2% |
| TM3 | CLR M223 - CGRP T9 | 31.2% | 41.5% |
| TM3 | CLR Q216 - CGRP H10 | 38.9% | 66.4% |
| TM3 | CLR Y227 - CGRP T6 | 55.9% | 32.7% |
| TM5 | CLR C299 - CGRP A5 | n/a | 32.0% |
| TM5 | CLR C299 - CGRP T4 | 38.3% | 50.5% |
| TM5 | CLR C299 - CGRP T6 | n/a | 36.1% |

| | | | |
|---|---|---|---|
| TM5 | CLR H295 - CGRP C2 | 30.1% | n/a |
| TM5 | CLR H295 - CGRP C7 | 95.4% | 88.7% |
| TM5 | CLR H295 - CGRP H10 | 63.5% | 52.1% |
| TM5 | CLR H295 - CGRP T6 | 94.4% | 87.3% |
| TM5 | CLR I298 - CGRP T6 | 41.8% | 48.3% |
| TM5 | CLR L291 - CGRP C2 | 37.9% | n/a |
| TM5 | CLR L291 - CGRP C7 | 31.2% | 33.0% |
| TM5 | CLR L291 - CGRP H10 | 40.2% | 48.1% |
| TM5 | CLR Y292 - CGRP A1 | n/a | 41.8% |
| TM5 | CLR Y292 - CGRP C2 | 71.6% | 79.0% |
| TM6 | CLR F349 - CGRP A5 | 61.9% | 78.1% |
| TM6 | CLR F349 - CGRP T4 | n/a | 50.8% |
| TM6 | CLR F349 - CGRP T6 | 45.4% | n/a |
| TM6 | CLR P353 - CGRP T4 | n/a | 32.0% |
| TM6 | CLR W354 - CGRP D3 | n/a | 46.0% |
| TM7 | CLR D366 - CGRP R11 | 99.1% | 91.3% |
| TM7 | CLR D366 - CGRP V8 | 58.9% | n/a |
| TM7 | CLR H370 - CGRP L12 | 78.3% | 73.7% |
| TM7 | CLR H370 - CGRP V8 | 72.3% | 79.2% |
| TM7 | CLR H374 - CGRP T9 | 44.3% | 47.5% |
| TM7 | CLR M369 - CGRP A5 | 65.9% | 49.6% |
| TM7 | CLR M369 - CGRP T4 | 34.0% | n/a |
| TM7 | CLR M369 - CGRP V8 | 41.0% | 52.5% |
| TM7 | CLR M373 - CGRP A5 | 31.4% | n/a |
| TM7 | CLR M373 - CGRP T6 | 41.0% | 32.5% |
| TM7 | CLR M373 - CGRP T9 | 43.2% | 67.1% |
| TM7 | CLR M373 - CGRP V8 | 52.8% | 50.6% |
| TM7 | CLR Y365 - CGRP D3 | 30.1% | n/a |

**Table S3.** Frequency of all polar and non-polar contacts occurred at the CLR-RAMP1 interface. Interactions occurred more than 40% of concatenated MD trajectories are presented. The interactions occurred less than 40% of total concatenated trajectories are not shown (n/a). ECD: Extracellular ectodomain, ECL2: Extracellular loop 2, TM3-4-5: Transmembrane domains 3-4-5.

| CLR Domain | Residue Pairs | Gs | Gs + CGRP | Rimegepant site1 | Rimegepant site2 | Telcagepant site1 | Telcagepant site2 |
|---|---|---|---|---|---|---|---|
| ECD | CLR D70 - RAMP1 P85 | 44.0% | 66.8% | 90.7% | n/a | 74.6% | n/a |
| ECD | CLR D90 - RAMP1 R112 | 40.9% | n/a | n/a | n/a | n/a | n/a |
| ECD | CLR E47 - RAMP1 G108 | 53.2% | n/a | n/a | 57.3% | n/a | n/a |
| ECD | CLR E47 - RAMP1 I106 | 44.5% | 49.7% | n/a | 67.7% | 55.7% | 51.5% |
| ECD | CLR E47 - RAMP1 R109 | 98.0% | 42.6% | 91.5% | 60.9% | 55.0% | n/a |
| ECD | CLR E47 - RAMP1 R112 | 49.5% | 56.6% | n/a | n/a | 44.0% | 50.4% |
| ECD | CLR E47 - RAMP1 S107 | 47.3% | n/a | n/a | 59.4% | 44.0% | 43.8% |
| ECD | CLR G71 - RAMP1 P85 | n/a | n/a | 52.8% | n/a | 53.9% | n/a |
| ECD | CLR I52 - RAMP1 L94 | 53.5% | 55.5% | 41.2% | 48.1% | 43.6% | 57.3% |
| ECD | CLR K51 - RAMP1 I106 | n/a | n/a | n/a | 43.9% | n/a | n/a |
| ECD | CLR M42 - RAMP1 A70 | 63.7% | 63.1% | 99.2% | 73.5% | 99.2% | 63.1% |
| ECD | CLR M42 - RAMP1 R67 | 75.6% | 77.7% | 58.7% | 67.5% | 51.5% | 74.9% |
| ECD | CLR M42 - RAMP1 W59 | 79.3% | 74.1% | 71.3% | 68.1% | 75.8% | 65.3% |
| ECD | CLR M42 - RAMP1 Y66 | 52.8% | 51.0% | 55.3% | 58.0% | 76.2% | 54.9% |
| ECD | CLR M53 - RAMP1 G98 | 56.7% | 58.4% | 53.0% | 68.4% | 48.1% | 56.7% |
| ECD | CLR M53 - RAMP1 H97 | 64.2% | 66.6% | 66.1% | 72.0% | 62.1% | 69.3% |
| ECD | CLR M53 - RAMP1 L94 | 62.5% | 56.2% | 56.3% | 52.8% | 43.2% | 50.7% |
| ECD | CLR M53 - RAMP1 R102 | 58.7% | 57.1% | 59.3% | 60.4% | 57.7% | 53.8% |
| ECD | CLR N39 - RAMP1 I63 | 79.4% | 75.9% | 77.2% | 82.8% | 69.4% | 69.6% |
| ECD | CLR N39 - RAMP1 R67 | n/a | 45.0% | 95.2% | 74.0% | 94.0% | 66.2% |
| ECD | CLR Q45 - RAMP1 F93 | 65.2% | 65.4% | 85.0% | 54.4% | 83.8% | |
| ECD | CLR Q45 - RAMP1 F93 | n/a | n/a | n/a | n/a | n/a | 45.2% |
| ECD | CLR Q45 - RAMP1 W84 | n/a | n/a | 87.6% | n/a | 86.8% | n/a |
| ECD | CLR Q45 - RAMP1 Y66 | 71.4% | 70.4% | 76.3% | 64.0% | 76.7% | 59.0% |
| ECD | CLR Q50 - RAMP1 C104 | 74.7% | 82.9% | 66.7% | 61.5% | 81.4% | 69.0% |
| ECD | CLR Q50 - RAMP1 F101 | 99.5% | 99.7% | 98.8% | 99.5% | 99.9% | 96.5% |
| ECD | CLR Q50 - RAMP1 H97 | 99.3% | 97.6% | 98.2% | 97.1% | 97.4% | 90.4% |
| ECD | CLR Q50 - RAMP1 I106 | 66.3% | 66.5% | 65.0% | 77.6% | 71.0% | 72.8% |
| ECD | CLR Q50 - RAMP1 P105 | n/a | n/a | n/a | 88.1% | 47.7% | 68.6% |
| ECD | CLR Q54 - RAMP1 I106 | n/a | 40.6% | 48.5% | 50.1% | 40.4% | 53.0% |
| ECD | CLR R119 - RAMP1 D90 | n/a | n/a | n/a | 47.5% | n/a | n/a |
| ECD | CLR R119 - RAMP1 F83 | 54.1% | 70.9% | 89.2% | n/a | 77.8% | 40.1% |
| ECD | CLR R119 - RAMP1 P85 | n/a | n/a | n/a | 44.8% | n/a | n/a |
| ECD | CLR R38 - RAMP1 D71 | 93.9% | 96.6% | 96.2% | 82.1% | 94.1% | 98.7% |
| ECD | CLR R38 - RAMP1 R67 | 60.0% | 63.9% | 70.8% | 52.8% | 54.7% | 68.8% |
| ECD | CLR S117 - RAMP1 F83 | n/a | n/a | n/a | 51.5% | n/a | n/a |

| | | | | | | | |
|---|---|---|---|---|---|---|---|
| ECD | CLR T43 - RAMP1 R109 | 75.7% | n/a | 45.9% | n/a | 43.2% | n/a |
| ECD | CLR T43 - RAMP1 W59 | 99.0% | 96.7% | 92.8% | 98.9% | 95.4% | 99.4% |
| ECD | CLR T68 - RAMP1 D90 | n/a | 54.7% | n/a | n/a | n/a | n/a |
| ECD | CLR W69 - RAMP1 P85 | n/a | n/a | 61.5% | n/a | 48.1% | n/a |
| ECD | CLR Y46 - RAMP1 F101 | 68.5% | 70.7% | 69.3% | 74.8% | 71.0% | 71.3% |
| ECD | CLR Y46 - RAMP1 G108 | n/a | n/a | n/a | 50.9% | n/a | n/a |
| ECD | CLR Y46 - RAMP1 H97 | 99.7% | 99.7% | 99.5% | 99.8% | 99.2% | 98.6% |
| ECD | CLR Y46 - RAMP1 R109 | n/a | n/a | n/a | n/a | n/a | 44.8% |
| ECD | CLR Y46 - RAMP1 W59 | 99.3% | 96.5% | 99.0% | 99.1% | 99.7% | 97.4% |
| ECD | CLR Y46 - RAMP1 Y66 | 70.9% | 76.1% | 68.3% | 66.3% | 60.2% | 64.3% |
| ECD | CLR Y49 - RAMP1 D90 | 97.1% | 98.0% | 99.8% | 92.1% | 96.2% | 88.0% |
| ECD | CLR Y49 - RAMP1 F93 | 92.8% | 96.6% | 93.5% | 88.4% | 89.7% | 90.0% |
| ECD | CLR Y49 - RAMP1 H97 | 86.3% | 86.7% | 84.4% | 91.7% | 90.5% | 92.0% |
| ECD | CLR Y49 - RAMP1 L94 | 42.7% | 52.2% | 45.8% | n/a | 52.3% | n/a |
| ECL2 | CLR D280 - RAMP1 R109 | n/a | n/a | n/a | 45.9% | n/a | n/a |
| ECL2 | CLR D280 - RAMP1 R112 | n/a | n/a | 97.5% | 42.5% | n/a | n/a |
| ECL2 | CLR D287 - RAMP1 R112 | 47.0% | n/a | 49.3% | n/a | n/a | n/a |
| ECL2 | CLR H289 - RAMP1 D113 | 99.6% | n/a | 98.8% | 90.4% | 59.7% | 86.4% |
| ECL2 | CLR H289 - RAMP1 I123 | n/a | 60.1% | n/a | n/a | n/a | n/a |
| ECL2 | CLR H289 - RAMP1 L119 | 72.7% | n/a | 62.2% | 68.3% | 61.8% | 55.0% |
| ECL2 | CLR H289 - RAMP1 Y120 | n/a | 45.2% | n/a | n/a | n/a | n/a |
| ECL2 | CLR L290 - RAMP1 D113 | n/a | n/a | n/a | 42.8% | n/a | n/a |
| ECL2 | CLR L290 - RAMP1 L119 | n/a | 58.7% | 41.1% | n/a | 45.9% | n/a |
| ECL2 | CLR L290 - RAMP1 P114 | 52.9% | n/a | 75.0% | n/a | n/a | 50.5% |
| ECL2 | CLR T288 - RAMP1 D113 | 97.1% | n/a | 91.6% | 86.8% | 52.3% | 40.5% |
| ECL2 | CLR T288 - RAMP1 R112 | n/a | n/a | 48.5% | n/a | n/a | n/a |
| ECL2 | CLR Y277 - RAMP1 A110 | n/a | n/a | n/a | n/a | 48.4% | n/a |
| ECL2 | CLR Y277 - RAMP1 D113 | n/a | n/a | 43.5% | n/a | n/a | n/a |
| ECL2 | CLR Y277 - RAMP1 G108 | 41.9% | n/a | | n/a | n/a | n/a |
| ECL2 | CLR Y277 - RAMP1 P114 | 78.1% | n/a | 82.9% | 75.8% | 76.5% | 77.5% |
| ECL2 | CLR Y277 - RAMP1 R112 | 45.4% | n/a | 68.3% | 51.5% | 40.4% | n/a |
| ECL2 | CLR Y277 - RAMP1 V111 | 51.4% | n/a | 41.0% | n/a | 64.2% | n/a |
| ECL2 | CLR Y278 - RAMP1 A110 | n/a | n/a | n/a | n/a | 55.2% | n/a |
| ECL2 | CLR Y278 - RAMP1 D113 | n/a | n/a | n/a | 58.7% | 45.7% | n/a |
| ECL2 | CLR Y278 - RAMP1 R112 | 49.2% | n/a | 94.5% | 58.5% | 51.6% | 72.4% |
| ECL2 | CLR Y278 - RAMP1 V111 | n/a | n/a | n/a | n/a | 76.4% | 62.8% |

| | | | | | | | |
|---|---|---|---|---|---|---|---|
| ICL2 | CLR E248 - RAMP1 R143 | n/a | n/a | 46.7% | n/a | n/a | n/a |
| ICL2 | CLR E248 - RAMP1 S141 | 56.5% | n/a | n/a | n/a | n/a | n/a |
| ICL2 | CLR F246 - RAMP1 S141 | n/a | n/a | n/a | n/a | 47.7% | n/a |
| ICL2 | CLR H251 - RAMP1 Q140 | 49.7% | 67.9% | n/a | n/a | 56.6% | n/a |
| ICL2 | CLR H251 - RAMP1 S141 | n/a | 63.9% | 67.3% | 59.4% | 42.6% | 50.4% |
| ICL2 | CLR V243 - RAMP1 K142 | n/a | 46.1% | n/a | n/a | n/a | n/a |
| TM3 | CLR F228 - RAMP1 T130 | 54.9% | 58.9% | 55.8% | 48.8% | 58.3% | 51.8% |
| TM3 | CLR I235 - RAMP1 S141 | 44.0% | n/a | n/a | n/a | 46.1% | n/a |
| TM3 | CLR I235 - RAMP1 V137 | n/a | 50.8% | 49.8% | 43.2% | 51.0% | 46.3% |
| TM3 | CLR I235 - RAMP1 V138 | 47.7% | n/a | n/a | n/a | 51.4% | n/a |
| TM3 | CLR L231 - RAMP1 T134 | 61.5% | 71.0% | 63.3% | 69.0% | 71.7% | 67.2% |
| TM3 | CLR T239 - RAMP1 S141 | 69.2% | 59.5% | 46.4% | | 49.6% | 40.0% |
| TM4 | CLR F262 - RAMP1 T130 | 85.4% | 72.5% | 75.0% | 80.8% | 83.8% | 82.5% |
| TM4 | CLR F262 - RAMP1 V129 | 45.6% | n/a | 42.0% | n/a | 54.7% | 43.8% |
| TM4 | CLR F262 - RAMP1 V133 | 45.0% | 41.5% | 50.6% | 56.7% | n/a | n/a |
| TM4 | CLR I269 - RAMP1 F122 | 69.2% | 40.0% | 47.8% | | 48.5% | 68.0% |
| TM4 | CLR L258 - RAMP1 V133 | 54.8% | 53.6% | 57.8% | 45.4% | 67.0% | 60.7% |
| TM4 | CLR W254 - RAMP1 Q140 | 84.5% | 59.1% | 87.8% | 86.8% | 87.8% | 76.5% |
| TM4 | CLR W254 - RAMP1 V137 | n/a | 47.7% | n/a | n/a | 41.7% | n/a |
| TM4 | CLR Y255 - RAMP1 S141 | 78.1% | 41.0% | n/a | n/a | 61.2% | n/a |
| TM4 | CLR Y255 - RAMP1 V137 | 42.4% | n/a | n/a | n/a | 53.4% | n/a |
| TM5 | CLR G296 - RAMP1 I127 | 75.0% | 53.0% | 69.7% | 74.1% | 71.8% | 76.6% |
| TM5 | CLR I293 - RAMP1 F122 | 63.1% | 47.9% | 65.1% | 66.2% | 58.2% | 63.4% |
| TM5 | CLR I293 - RAMP1 I123 | n/a | 42.5% | n/a | 46.4% | n/a | n/a |
| TM5 | CLR I293 - RAMP1 P126 | n/a | 50.6% | n/a | n/a | n/a | n/a |
| TM5 | CLR P297 - RAMP1 I127 | n/a | 40.0% | n/a | n/a | 51.9% | 58.5% |
| TM5 | CLR P297 - RAMP1 P126 | 41.7% | 43.1% | 40.0% | n/a | 48.1% | 50.8% |
| TM5 | CLR P297 - RAMP1 T130 | 72.1% | 75.5% | 72.7% | 77.7% | 64.9% | 63.0% |
| TM5 | CLR V304 - RAMP1 L131 | 45.3% | 53.1% | 40.8% | 52.2% | n/a | 40.2% |
| TM5 | CLR Y292 - RAMP1 I123 | 54.3% | 51.7% | 63.7% | 56.7% | 58.8% | 61.5% |

**Table S4.** Prepared systems for Molecular Dynamics Simulations

| **CLR with Gs** | **CLR without Gs** |
|---|---|
| 1- CLR-CGRP-Gs | 5- CLR-RAMP1-rimegepant-site1 |
| 2- CLR-Gs | 6- CLR-RAMP1-rimegepant-site2 |
| 3- CLR-RAMP1-CGRP-Gs | 7- CLR-RAMP1-telcagepant-site1 |
| 4- CLR-RAMP1-Gs | 8- CLR-RAMP1-telcagepant-site2 |
| | 9- CLR-rimegepant-site2 |
| | 10- CLR-telcagepant-site2 |

**Table S5.** CLR topology in UNIPROT vs. our study according to the POPC membrane and protein orientation

| Topology | Position interval in UNIPROT | Position interval in our study |
|---|---|---|
| Extracellular | 23 – 146 | 33 - 136 |
| TM1 | 147 – 166 | 137 - 166 |
| ICL1 | 167 – 173 | 167 - 173 |
| TM2 | 174 – 193 | 174 - 197 |
| ECL1 | 194 – 213 | 198 - 212 |
| TM3 | 214 – 236 | 213 - 240 |
| ICL2 | 237 – 253 | 241 - 252 |
| TM4 | 254 – 273 | 253 - 275 |
| ECL2 | 274 – 289 | 276 - 290 |
| TM5 | 290 – 313 | 291 - 317 |
| ICL3 | 314 – 336 | 318 - 330 |
| TM6 | 337 – 354 | 331 - 354 |
| ECL3 | 355 – 366 | 355 - 360 |
| TM7 | 367 – 388 | 361 - 388 |
| Cytoplasmic helix - VIII | 389 – 461 | 389 - 418 |
